# Site-Specific Entry Factors Define Cellular Susceptibility to SARS-CoV-2 in Human Tissues

**DOI:** 10.64898/2026.05.07.723425

**Authors:** Mireia Vivas Roca, Cristina Mancebo-Pérez, Mohmed Abdalfttah, Ismael Balafkir, Aleix Benítez-Martínez, Víctor Montal, Alba González, Irene Mota-Gómez, Judith Grau-Expósito, Daan KJ Pieren, Bibiana Planas, Joel Rosado, María Eugenia Semidey, Stefania Landolfi, Eloy Espín-Bassany, Enric Trilla, Vicenç Falcó, Adrià Curran, Joaquín Burgos, María J Buzon, Víctor Guallar, Holger Heyn, Meritxell Genescà

## Abstract

SARS-CoV-2 primarily infects the respiratory epithelium through entry factors ACE2 and TMPRSS2. However, alternative viral entry receptors and/or modulators of viral entry factors expression may contribute to infection in respiratory as well as non-respiratory tissues, including the kidneys and colon. Using single-cell analyses, we identified novel candidates for spike-mediated viral entry in epithelial cells from diverse fresh human tissues exposed to a labeled spike-pseudotyped virus. Systematic viral tracking revealed tissue-specific membrane receptors that contribute to viral entry, functionally validated in human tissues and partially predicted by molecular modeling. We found ADAM17 and IL1R1, together with canonical molecules, to significantly facilitate viral entry in the lung parenchyma, while entry into the renal cortex was less dependable on canonical factors, but strongly modulated by JAK1/2 engagement. These findings uncover distinct viral entry mechanisms across tissues and suggest novel organ-specific therapeutic targets for patients at risk.

## INTRODUCTION

Severe acute respiratory syndrome coronavirus 2 (SARS-CoV-2), the etiological agent of COVID-19, displays broad tissue tropism that extends beyond the respiratory tract and underlies the systemic manifestations of disease. In addition to infection of the upper and lower airways, viral RNA, protein, and associated pathology have been detected in the gastrointestinal tract, kidney, heart, and central nervous system, implicating direct viral entry into multiple organ systems in disease pathogenesis ^1–4^. The molecular mechanisms that govern SARS-CoV-2 entry into distinct human tissues, however, remain incompletely defined.

Viral entry is initiated by engagement of the trimeric spike (S) glycoprotein with host cell receptors, followed by proteolytic activation that enables membrane fusion. Angiotensin-converting enzyme 2 (ACE2) is the best-characterized SARS-CoV-2 receptor and is sufficient to confer susceptibility when ectopically expressed^5^. Viral entry was also found to depend on transmembrane protease serine 2 (TMPRSS2) or endo/lysosomal cathepsins^5,6^. Consistent with this, inhibition of TMPRSS2 markedly reduces viral entry in airway epithelial cells and lung-derived primary cells^5,7^. ACE2 is co-expressed with TMPRSS2 in discrete epithelial populations of the nasal mucosa, airways, and alveoli^3,8–12^, as well as in other tissues, including gut enterocytes^13^. In the respiratory tract, ciliated cells, followed by club cells within the airways, and alveolar type 2 (AT2) cells lining the alveoli, are among the cell types with the highest level of ACE2 and TMPRSS2 expression^8,10–12,14^

However, additional factors are involved in viral entry into the host cell, which collectively influence entry efficiency^15,16^. Cofactors and alternative receptors may facilitate virus-host cell interactions in cells with low ACE2 expression^17^. Indeed, metalloproteases including ADAM10 and ADAM17 further modulate viral entry by regulating ACE2 shedding and spike processing, thereby influencing receptor availability and fusion efficiency^18–21^. SARS-CoV-2 can also utilize endosomal entry routes that depend on cathepsins, as well as proprotein convertases such as furin, which cleaves spike at the S1/S2 junction and enhances infectivity^11,22^. Additionally, several non-canonical receptors and attachment factors, including CD147 (BSG), AXL, ASGR1, and KREMEN1, have been proposed to facilitate SARS-CoV-2 binding or internalization under specific conditions^19,23,24^. Recent work has further expanded the repertoire of SARS-CoV-2 entry mechanisms by identifying TMEM106B as an ACE2-independent spike receptor capable of directly engaging the receptor-binding domain and mediating viral entry and syncytium formation^25^. These findings underscore the plasticity of SARS-CoV-2 entry pathways and suggest that distinct tissues may rely on different combinations of canonical and alternative host factors to support infection.

Advances in high-throughput and single-cell transcriptomics have provided valuable insights into the cell-type–specific expression of SARS-CoV-2 entry genes across human tissues. A comprehensive single-cell RNA sequencing (scRNA-seq) meta-analysis demonstrated that ACE2, TMPRSS2, and CTSL are co-expressed in specific nasal, airway, and alveolar epithelial populations^11^, although protein expression under physiological conditions may be low in some of these populations. Complementing this, Singh et al. ^26^ analyzed a panel of 28 viral receptors and host factors, revealing potential non-canonical entry pathways in the lung and highlighting the nasal epithelium as a major hub of both pro-viral and antiviral factor expression. Newer functional genomics efforts have used CRISPR-based screening combined with single-cell readouts to systematically test hundreds of candidate host factors for their effect on viral infection dynamics and host transcriptional response, underscoring genes such as EIF4E2, EIF4H, NFKBIA^27^, PLSCR1^28^ and components of V-ATPases, ESCRT, and N-glycosylation pathways ^29^ as critical host dependency factors influencing viral replication efficiency and host response. Furthermore, molecular docking has emerged as a valuable computational strategy for identifying and refining viral factors, such as NRP1 as an attachment/co-factor that potentiates ACE2-dependent entry^17,30–32^. Still, their discovery and validation has frequently relied on cell lines^33^, which lack the complexity of human tissues^34^. More recently, human airway, renal and intestinal organoids, which partially replicate physiological conditions and individual diversity^35^, have provided crucial evidence for the functional roles of ACE2, TMPRSS2, and additional host factors in SARS-CoV-2 entry^4,36–39^.

Despite these discoveries, the observed heterogeneity in tissue tropism and inter-patient variability remains incompletely explained, underscoring the need for a more comprehensive examination of virus-cell interactions across diverse human tissues. To address these gaps, a systematic and functional interrogation of SARS-CoV-2 entry across primary human tissues is required. In this study, we combined single-cell analyses with an experimental platform in which sorted epithelial cells from human nasal, lung and renal cortex samples were exposed to the same labeled spike-pseudotyped virus. This approach enabled a direct, quantitative comparison of viral entry across tissues under controlled conditions. By integrating this data with molecular modeling predictions and functional inhibition assays in intact human tissues, we identified tissue-specific membrane receptors and modulators that contribute to viral entry. Together, these approaches revealed distinct entry patterns across tissues and individuals. In cells from lung parenchyma, viral entry is strongly dependent on ACE-2, TMPRSS2 and ADAM17, but we also underscore a role for IL1R1. In contrast to the lung, viral entry into the renal cortex is less dependable on ACE-2 and TMPRSS2 factors and remarkably involves the JAK/STAT pathway. Inhibition of this pathway is as efficient as blocking the canonical receptors. Importantly, we reveal the relative contribution of a broader range of receptors to viral entry susceptibility across human tissues. Our findings provide a more holistic “atlas” of permissivity in SARS-CoV-2 cellular targets derived from human tissues and highlight the complexity of SARS-CoV-2 entry in clinical context.

## RESULTS

### Isolation of epithelial cells permissive to SARS-CoV-2 entry in multiple tissues and phenotypic characterization

To dissect epithelial susceptibility to SARS-CoV-2 across human tissues, we *ex vivo* infected cell suspensions obtained from nasal brush pools, lung parenchyma, renal cortex and colon biopsies with a labeled SARS-CoV-2-GFP^+^ pseudovirus based on Vesicular Stomatitis Virus (VSV) expressing the D614G spike (S; VSV-GFP-S). Nasal samples were obtained from pooled nasal brushes of 7-10 healthy volunteers between 23-65 years old^40^. Non-neoplastic lung parenchyma, kidney and colon biopsies were obtained from patients undergoing resection at Hospital Universitari Vall d’Hebron (HUVH). Infection and digestion protocols were individually optimized to ensure detection of infection and high cellular viability across tissues for subsequent sorting. We designed specific spectral flow cytometry panels for each tissue, based on predicted susceptible cellular subsets, at the time that the study was initiated. Nasal mucosa and lung parenchyma phenotyping of CD45^-^CD31^-^EpCAM^+^ epithelial cells, included integrin subunit alpha 6 (ITGA6) and nerve growth factor receptor (NGFR) for basal cells, acetylated α-tubulin enriched in ciliated cells, Mucin 5AC (MUC5AC) for goblet cells, and surfactant Protein-C (SPC), HLA-DR (DR) and EpCAM for alveolar type-II (AT-II) ^3,7,12,41–43^. Kidney epithelial subsets were characterized by EpCAM and E-cadherin expression, CD10 together with CD13 for proximal tubule epithelial cells (PTEC) and podocalyxin for podocytes^44^.

Limited cell yields from nasal brushings precluded inclusion of a negative control for background. Still, out of EpCAM^+^ cells obtained from three large pools of nasal samples, ITGA6 (57.30% [46.20-57.50], ***Figure 1A-C***) and high HLA-DR (45.7% [43.40-46.50]) were most frequently expressed, followed by NGFR^+^ and double ITGA6^+^NGFR^+^ (38.30% [23.90-40.60] and 30.60% [13.38-32.90], respectively), which expression may indicate basal or basal stem cell phenotype^43^. Of note, expression of α-tubulin^+^ and MUC5AC^+^, typically expressed in ciliated and goblet cells respectively^43^, was very low (***Figure 1A-C***), which may be due to sampling site, brushing intensity or sample processing^45,46^. Among GFP⁺ cells, which represented a median of 0.83% [0.834-1.830] of total EpCAM^+^ cells, ITGA6 was expressed in nearly 90% of them (88.10% [80.20-92.30], ***Figure 1B***), frequently together with HLA-DR or NGFR (***Figure 1D***). In fact, NGFR^+^ and ITGA6^+^NGFR^+^ expression was enriched among EpCAM⁺GFP⁺ cells compared to the total EpCAM⁺ population (***Figure 1C-D***). Additional unsupervised clustering analyses based on mean fluorescence intensity identified 8 metaclusters (MC) similarly expressed in the three pools of samples (***Figure S1A***), where MC#7 and #8 included the highest GFP expression (***Figure S1B***). These analyses confirmed the high expression of ITGA6 in these GFP^+^ clusters, together with α-tubulin enrichment for #7 or with NGFR for #8 (***Figure S1C***).

**Figure 1.**
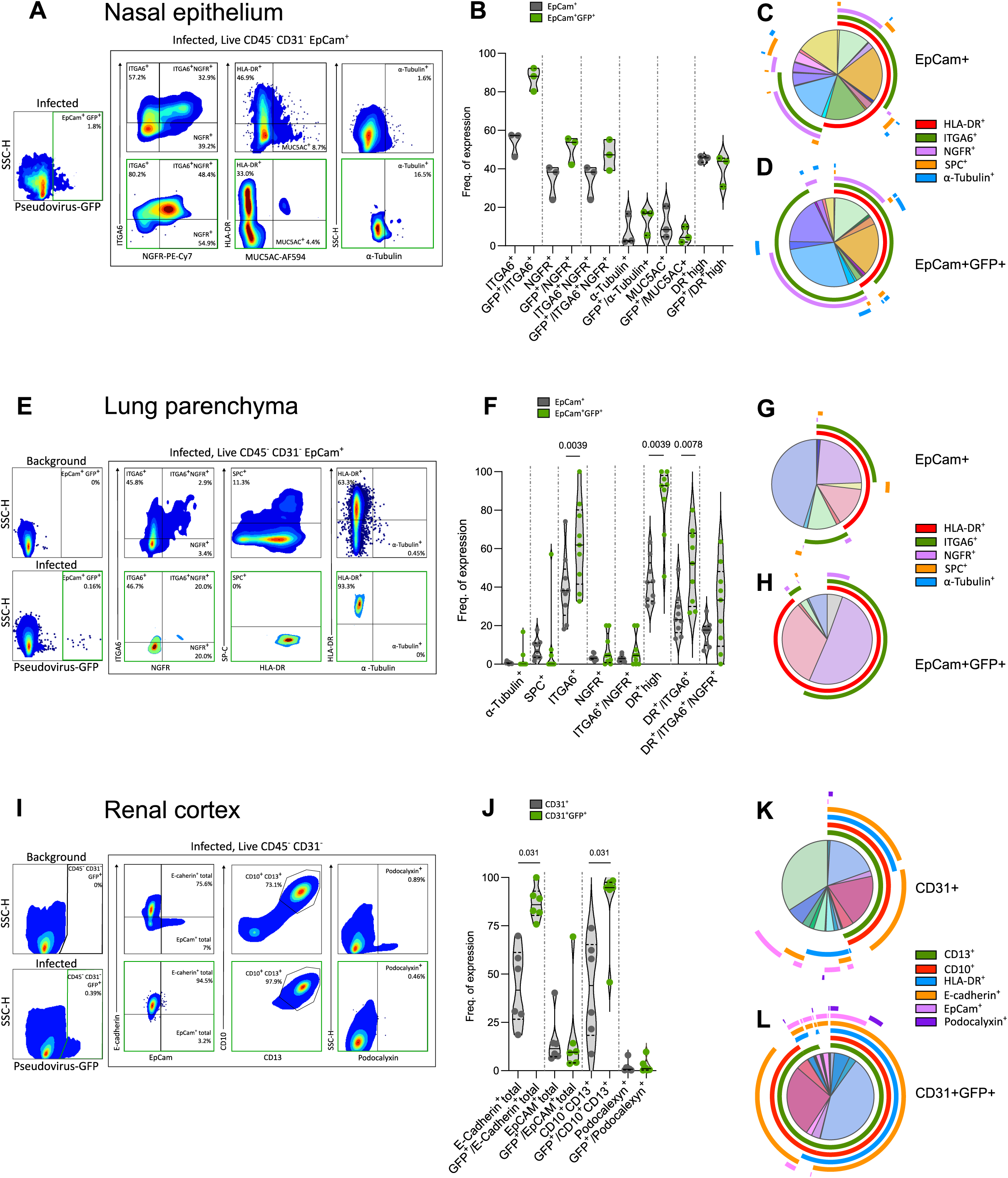
Isolation of epithelial cells permissive to SARS-CoV-2 entry in multiple tissues and phenotypic characterization. (A-D) SARS-CoV-2 pseudovirus *ex vivo* infected nasal epithelial cells (n=3 pools). **(E-H)** SARS-CoV-2 pseudovirus *ex vivo* infected lung epithelial cells (n= 9). **(I-L)** SARS-CoV-2 pseudovirus *ex vivo* infected renal cortex cells (n=7). Flow cytometry plots showing the phenotypic comparison between GFP⁺ (infected, green) and GFP⁻ (uninfected) cells of CD45^-^CD31^-^EpCAM^+^ cells from the **(A)** pool #NAL02 (n= 7) or **(E)** #HLTE197, which includes a paired sample exposed to a spike-empty pseudovirus (background). **(B and F)** Violin plots depicting the frequency (%) of different epithelial marker expressions within total nasal (**B**) or pulmonary (**F**) EpCAM^+^ (grey) and EpCAM^+^GFP^+^ cells (green). **(C-D and G-H)** Boolean pie charts displaying the proportion of nasal (**C-D**) or pulmonary (**G-H**) EpCAM^+^ (**C-G**) and EpCAM^+^GFP^+^ (**D-H**) cells expressing combinations of color code molecules according to the legend and indicated as surrounding arcs around the pie chart. **(I)** Flow cytometry plots showing the phenotypic comparison of CD45^-^CD31^-^ cells from #RINN22, either infected (GFP⁺; green) or exposed to a ‘background’ pseudovirus. **(J)** Violin plots depicting the frequency (%) of various molecules within total CD31^-^ (grey) and CD31^-^GFP^+^ cells (green). **(K-L)** Boolean pie charts displaying the proportion of CD31^-^ (**K**) and CD31^-^GFP^+^ (**L**) cells expressing combinations of color code molecules according to the legend and indicated as surrounding arcs around the pie chart. For all violin plots, data are represented as median ± IQR. Statistical analyses were performed using two-sided nonparametric Wilcoxon matched-pairs signed-rank test.

Similarly, in the lung parenchyma, regardless of high sample variability, we detected abundant expression of ITGA6 and HLA-DR in epithelial cells (38.20% [25.35-49.35] and 42.50% [32.80-52.65] respectively, ***Figure 1E-F***), which were co-expressed in 23.10% [16.40-31.85] of these cells (***Figure 1E-F***). SPC^+^ cells were detected in low frequency, followed by also scarce NGFR^+^ and α-tubulin^+^ epithelial cells (***Figure 1F-G*)**. The most permissive lung cells were HLA-DR^high^ cells, as reported before^7^, but also ITGA6^+^, with frequent co-expression of both proteins (***Figure 1H***). In fact, all these three phenotypes, single HLA-DR^high^ or ITGA6^+^, and double HLA-DR^high^ITGA6^+^ were significantly enriched in epithelial GFP+ cells, which represented 0.18% [0.10-0.43], compared to the total epithelial fraction (p = 0.0039 for single expression and p = 0.0078 for double, ***Figure 1F-H***), followed by HLA-DR^high^ITGA6^+^ subset (p=0.0078, ***Figure 1F***). Although ITGA6 expression is typically associated with airway/bronchial basal cells, AT2 cells, which can be identified as EpCAM^+^HLA-DR^+^ phenotype^7^, can also express intermediate ITGA6 levels^47,48^. Additional unsupervised clustering analyses confirmed these results, where the only GFP^+^ cluster was #7, that showed high HLA-DR expression and low ITGA6 (***Figure S2***).

For renal samples, we focused on the non-hematopoietic, non-endothelial fraction (determined by CD45^-^CD31^-^, ***Figure 1I-J***) to assess expression of the molecules of interest. In this way, the most frequent phenotypes, E-cadherin^+^ cells and CD10^+^CD13^+^ cells, were significantly enriched within the GFP^+^ permissive cell fraction (p = 0.031 for both, ***Figure 1J***). Although the frequency was variable, these differences were independent of EpCAM expression, which was in general low (***Figure 1J-K***). These results align with low and variable expression of EpCAM expression in proximal tubular epithelial along the nephron^49^. Among renal GFP⁺ cells (0.455% [0.030-0.7570]), CD10/CD13 and E-cadherin were co-expressed in ∼80% of them, together with HLA-DR in more than 50% of all GFP⁺ cells (***Figure 1L***), and these results were validated by unsupervised clustering analyses (***Figure S3***). We identified 17 clusters, from which #05 was identified as GFP^+^ CD10^+^CD13^+^ differentially expressed in infected compared to uninfected samples (***Figure S3***), indicating PTEC as the main target.

For colon samples, despite optimizing *ex vivo* infection with SARS-CoV-2 pseudoviral infection as performed with the rest of the tissues, detectable infection remained inconsistent and inefficient, preventing the recovery of sufficient GFP⁺ cells for high-quality single-cell RNA sequencing (***Figure S4A***). To address this limitation, we implemented a luciferase-based assay, as a more sensitive and scalable method to evaluate infection. With this approach, we detected infection in 61.5% of the colons tested, with positive cases predominantly localized in samples obtained from sigmoid region (***Figure S4B***).

Overall, SARS-CoV-2 spike-mediated entry in cells from our models varies by tissue type. In the nasal epithelium, despite cellular diversity, infected cells are EpCAM^+^ and enriched for ITGA6 together with NGFR or HLA-DR. In the lung, cells consistently express ITGA6 with or without HLA-DR expression. In the renal cortex, cells are predominantly E-cadherin⁺, CD10⁺, and CD13⁺ and are frequently HLA-DR⁺, while generally lacking EpCAM expression. *Ex vivo* pseudoinfection of colon samples was largely inefficient, with only regions of the sigmoid mucosa showing increased susceptibility.

### Tissue-resolved profiling of epithelial cell susceptibility to SARS-CoV-2 entry

To identify molecules influencing epithelial susceptibility to SARS-CoV-2 across human tissues, we applied single-cell RNA sequencing (Smart-seq2) on the purified epithelial cells from nasal, lung, and renal cortex, following *ex vivo* infection with the SARS-CoV-2 pseudovirus (***Figure 2***)^50^. GFP expression enabled discrimination between infected (GFP⁺) and uninfected (GFP⁻) cells, according to gating strategies indicated for each tissue (***Figure 1***), which were individually sorted for full-length single-cell transcriptome profiling. After quality control and normalization, unsupervised clustering and transcriptional analysis were performed to resolve cellular clusters associated with viral entry across tissues (***Figure 2A***).

**Figure 2.**
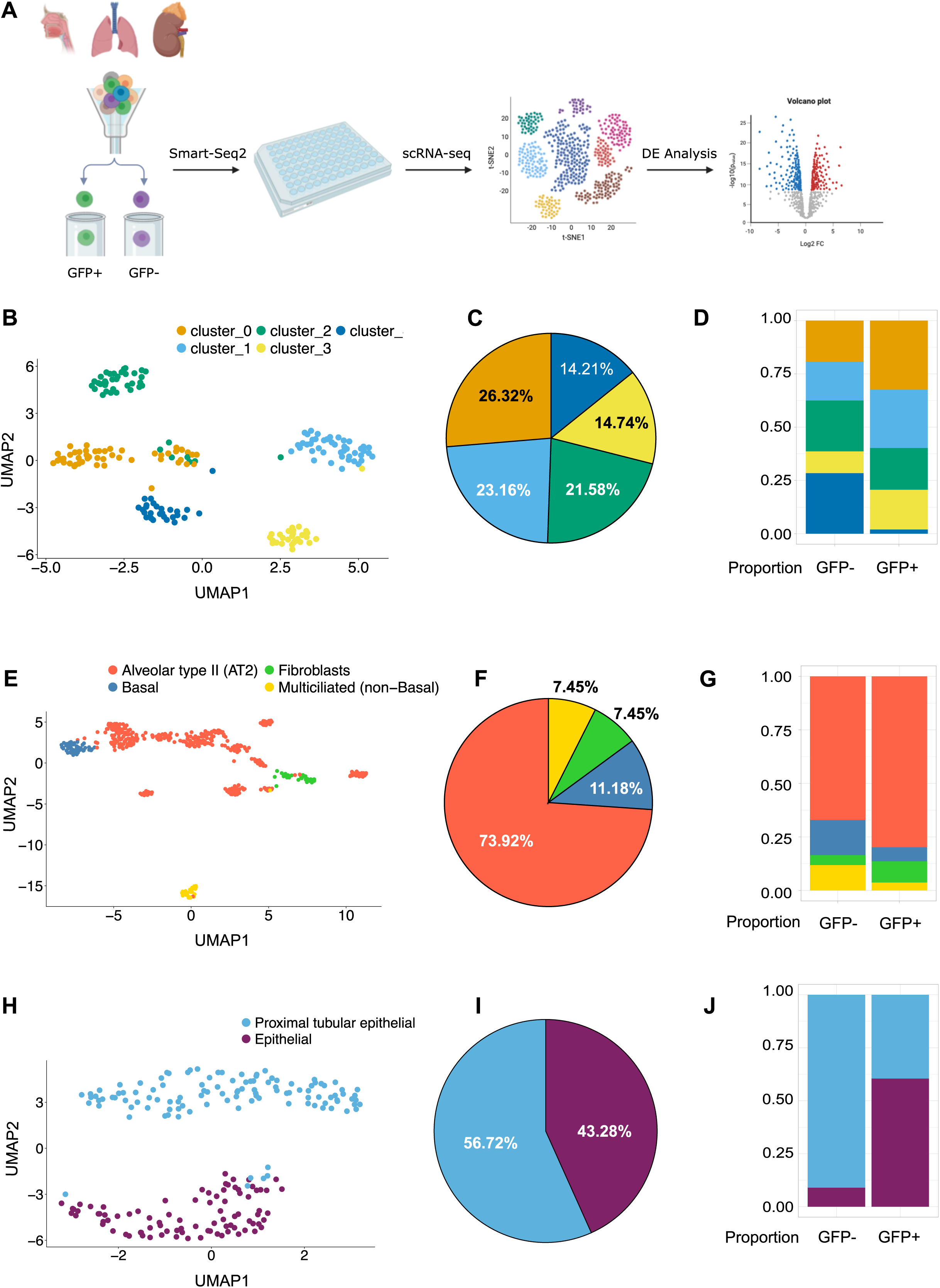
Transcriptional profiling of epithelial cell states following SARS-CoV-2 pseudovirus exposure across human tissues. **(A)** Schematic overview of the experimental workflow. Epithelial cells were isolated from nasal, lung, and kidney tissues and infected *ex vivo* with SARS-CoV-2 pseudovirus. GFP⁺ (infected) and GFP⁻ (uninfected) cells were sorted and processed using Smart-seq2. After quality control, cells were clustered and analyzed by tissue. **(B–D)** Nasal epithelial cells. **(B)** UMAP embedding of five transcriptionally distinct clusters. **(C)** Cluster distribution across all nasal epithelial cells. **(D)** Cluster proportions stratified by GFP⁺ and GFP⁻ conditions. **(E–G)** Lung epithelial cells. **(E)** UMAP embedding of lung-derived cells showing the separation of three epithelial and one stromal population. **(F)** Cluster distribution across epithelial and stromal populations. **(G)** Cluster proportions by infection status. **(H–J)** Renal cortex epithelial cells. **(H)** UMAP embedding of two epithelial clusters. **(I)** Cluster distribution across all renal epithelial cells. **(J)** Cluster proportions stratified by GFP⁺ and GFP⁻ conditions.

In the nasal compartment, epithelial cells were defined by the expression of canonical markers including *EPCAM, ELF3, CLDN4,* and *CDH1* (***Figure S5C***). Clustering analysis resolved five transcriptionally distinct populations (***Figure 2B***), that we could not precisely annotate according to previous publications^12^ (***Figure S5D***). These clusters were relatively balanced in abundance, with no single state dominating the nasal epithelium (***Figure 2C***). Notably, upon infection, cluster 0, 1 and 3 were enriched within the GFP⁺ fraction compared to GFP⁻ cells, while cluster 4 was not represented among GFP⁺ cells (***Figure 2D***). In the lung, epithelial cells were similarly identified by expression of *EPCAM* or *DCN*, which identified epithelial and stromal cells (***Figure S6***). As expected, epithelial cells constituted most lung-derived cells, however a small proportion of fibroblast (cluster 3) was detected, likely due to incomplete exclusion during the sorting process (***Figure 2E and S6B***). A total of four transcriptionally distinct states (***Figure 2E***) were identified: multiciliated epithelial cells, distinguished by *C9orf194* and *RSPH1;* AT2 cells, characterized by *SFTPC, SFTPB,* and *SFTPA;* and basal cells, characterized by *KRT15* and *KRT17*, among other genes (***Figure S6C***)^51^. The relative proportions of these states remained broadly stable across GFP⁺ and GFP⁻ fractions, although AT2 were dominant within the GFP fraction (80% of the cells), followed by fibroblast, basal and multicilliated cells; these last two subsets more abundant among GFP⁻ cells (***Figure 2G***), suggesting a reduced susceptibility to viral entry.

In renal cortex, epithelial cells were defined by *EPCAM* together with proximal tubular markers such as *AQP1* and *CUBN,* and distal tubules (***Figure S7***)^34,52^. Clustering analysis identified two major epithelial states: proximal tubular epithelial cells (PTEC) and a broader epithelial population (***Figure 2H***). PTEC constituted the dominant fraction of the compartment (***Figure 2I***), consistent with their prevalence in renal cortex. Yet, when comparing viral entry status, GFP⁺ cells were more frequently assigned to the undefined epithelial population, raising the possibility that this state may be more permissive to viral infection (***Figure 2J***). Overall, transcriptional analysis confirmed and further refined data revealed by phenotyping, identifying multiple clusters permissive to viral entry in nasal samples, AT2 cluster as the most permissive one in the lung parenchyma and epithelial cells, including PTEC for the renal cortex samples.

### Differential expression analysis identifies SARS-CoV-2 entry factors across human epithelial tissues

To identify candidate SARS-CoV-2 entry factors, we performed differential expression analyses (DEA) between GFP⁺ (infected) and GFP⁻ (uninfected) cells within each epithelial cluster of each tissue. We mostly focused on target proteins predicted to be expressed in the membrane of cells that were GFP⁺ as candidates to be involved in SARS-CoV-2 viral entry. Among the top upregulated genes across all three tissues, considering all clusters, we identified those involved in immune activation, including components of the interferon (IFN) response and innate antiviral signaling such as *NFKBIA, IFIT2, IFIT3* and *OASL* (***Figure 3A-C*** *and **Table S2***), confirming virus-induced immune activation and IFN signaling in GFP^+^ cells. Most of these molecules were discarded, as these were expected to be restricting factors less likely to participate in viral entry^1,27^.

**Figure 3.**
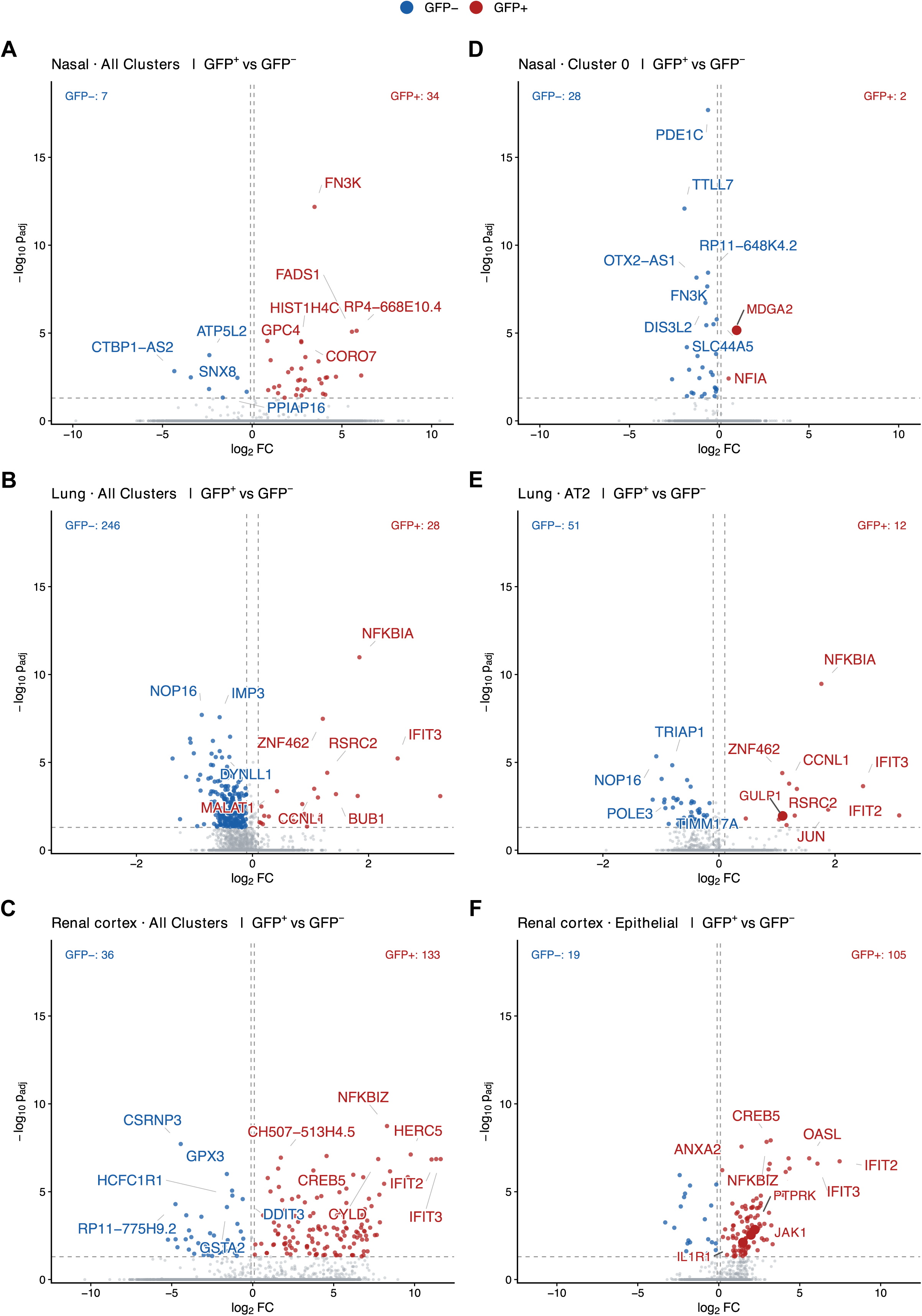
Differential Gene Expression Between GFP⁺ and GFP⁻ Populations Across Human Tissues. Volcano plots showing differentially expressed genes between GFP⁺ and GFP⁻ in overall epithelial cells from the **(A)** nasal mucosa, **(B)** lung parenchyma and **(C)** renal cortex; or in individual clusters from a given tissue: **(D)** epithelial cluster 0 from the nasal mucosa; **(F)** alveolar type 2 (AT2) cells from the lung; and **(G)** general epithelial cluster from renal cortex. The horizontal axis shows log₂ fold change, and the vertical axis shows –log₁₀ adjusted p values. Selected significantly upregulated genes (in red) are highlighted in bigger dots.

In the nasal epithelium, among all the clusters, only one candidate gene was detected in cluster 0, which was *MDGA2* with an average log₂FC of 0.54 and and p_adj_ of 3.7×10⁻⁵ (***Figure 3B*** and ***Table S2)***. MDGA2, a MAM domain-containing GPI-anchored protein, is known for its tumor suppressor role in gastric cancer and its function as a synaptic repressor^53^, unrelated to viral entry. In contrast, lung-derived cells revealed three candidate genes, *GULP1*, *ADAMTSL3*, and *PILRA,* across epithelial GFP^+^ subsets. *GULP1*, a PTB domain–containing adaptor protein involved in phagocytosis and apoptotic cell clearance ^54^, was enriched in GFP^+^ cells of the AT2 cluster (avg_log_2_FC = 1.06, p_adj_ = 0.012; ***Figure 3E and Table S2***). *ADAMTSL3*, a secreted matricellular protein from the ADAMTS-like family that has been associated with cardiac dysfunction^55^, was over-expressed in multiciliated GFP^+^ cells (avg_log2FC = 0.257, p_adj_ = 0.013; ***Table S2***). Also, paired immunoglobin like type 2 receptor alpha (*PILRA*), an immune regulator and entry coreceptor for Herpes simplex virus-1 (HSV-1) via glycoprotein B interaction^56^, was detected to be enriched in basal respiratory cells (avg_log_2_FC = 0.35, p_adj_ = 0.0001; ***Table S2***). Although not statistically significant, we included ADAM17 for further validation, a transmembrane metalloprotease regulating cell signaling via ectodomain shedding, that has been proposed to facilitate SARS-CoV-2 cell entry^18,20^ (weakly detected in AT2 cells; avg_log_2_FC = 0.234, p_adj_ > 0.05).

From renal clusters, several additional candidates were identified in the general epithelial cluster (***Figure 3F and Table S2***): cell adhesion molecule 1 (*CADM1*), a member of the immunoglobulin superfamily encoding a single cell-membrane glycoprotein primarily mediating intercellular adhesion (avg_log2FC = 2.60, p_adj_ = 0.000; ***Figure 3F***); interleukin 1 receptor, type I (*IL1R1*, avg_log2FC = 1.49, p_adj_ = 0.007; ***Figure 3F***) and Janus Kinase 1 gene (*JAK1*) (avg_log2FC = 1.99, p_adj_ = 0.003; ***Table S2***), mediators of inflammatory responses and immunological function; and protein tyrosine phosphatase receptor type K (*PTPRK*), a regulator of cell adhesion signaling (avg_log2FC = 2.23, p_adj_ = 0.001; ***Table S2***).

In addition to the candidates directly identified through scRNA-seq analysis, we specifically analyzed gene expression of previously identified molecules involved in SARS-CoV-2 viral entry, as well as genes used for phenotyping (***Table S3***). Although presenting very low mean gene expression in our tissue clusters, including ACE2 and TMPRRS2, the ratios between GFP^+^ over GFP^-^ captured the overexpression of these genes in GFP^+^ cells from individual tissue-clusters (***Table S4***). Of note, mean gene expression differences between GFP^+^ over GFP^-^ cells were not significant and, thus, not detected by DEA due to very low or inconsistent expression within clusters.

Altogether, nine candidates, including six new host proteins namely MDGA2 (from nasal), ADAMTSL3, GULP1, PILRα (from lung), and CADM1, PTPRK (from kidney), together with two proteins associated with the inflammatory response, IL1R1 and JAK1 (from kidney), and the already reported ADAM17^18,20^ (from lung), were selected for further analyses. Importantly, inhibitor molecules that could be used to block their function experimentally have been described for all of them.

### Functional validation of selected SARS-CoV-2 targets

Functional validation of selected candidates was performed on freshly isolated tissue-derived cells, as performed for single cell sorting. Considering samples availability and tissue size limitation, pre-screening to determine inhibitor concentration was limited to three doses and three samples for most cases (***Figure S8***). Of note, cell viability was assessed for all candidates in both lung and kidney, confirming the absence of cytotoxic effects (***Figure S8***). Due to the limitation of obtaining sufficient cell yield of viable nasal brush samples, we did not validate the panel of candidates in these samples.

As a reference control, we included inhibition of the canonical entry factors ACE2 and TMPRSS2 using an anti-ACE2 antibody and Camostat, as described previously^7^. As expected, in lung derived cells, anti-ACE2 reduced infection by a median of 96% [87–98], while Camostat, decreased infection by 93% [81–96] (***Figure 4A***). Additionally, inhibition of ADAM17 using KP-457 also significantly reduced viral entry by 77% [67–85] (***Figure 4A***). Interestingly, IL1R1 inhibition, using AF12198, was also capable of lowering viral entry by 60% [45–74], highlighting the contribution of inflammatory signaling modulating viral entry in target lung cells. In contrast, JAK1/2 inhibition using Ruxolitinib, which blocks JAK-STAT signalling, had no detectable effect. Among the other tested candidates, anti-ADAMTSL3, anti-CADM1 and anti-PILRα exhibited highly variable effects, reducing infection by more than 70% in some samples while showing no detectable effect in others (***Figure 4A***). The inhibitory effects of anti-GULP1, anti-MDGA2, and anti-PTPRK molecules also showed substantial variability across tissue samples, however with overall modest effects (***Figure 4A***).

**Figure 4.**
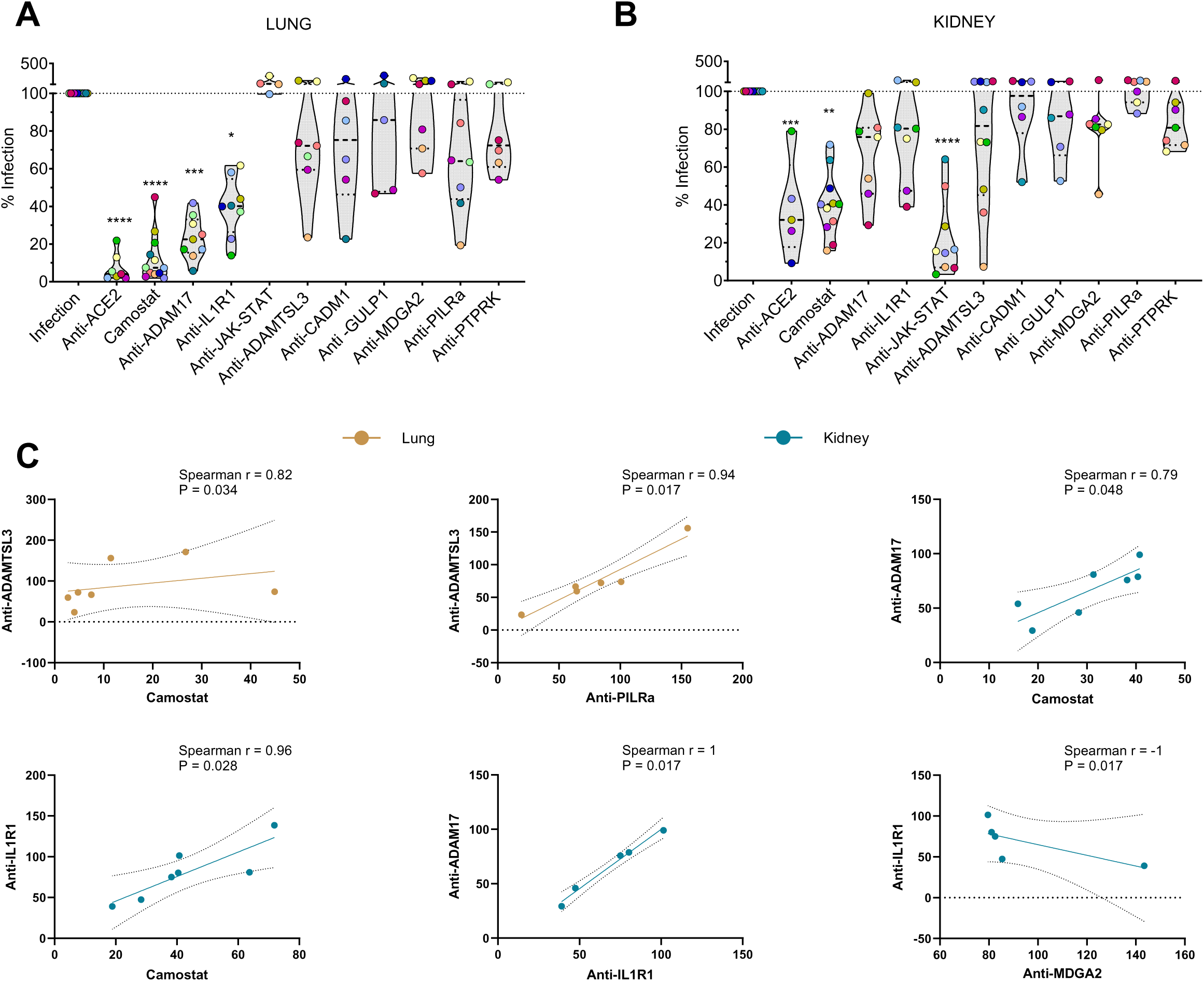
Functional validation of candidate entry factors in lung and renal tissue. (**A–B**) SARS-CoV-2 pseudovirus entry in lung (**A**) and kidney (**B**) epithelial cells following treatment with inhibitors. Lung or renal cortex-derived cells were exposed to pseudovirus in the presence of anti-ACE2 (25 µg/ml), Camostat (100 µM), KP-457 (ADAM17 inhibitor, 100 µM), anti-IL1R1 (100 µM), Ruxolitinib (JAK1/2 inhibitor, 100 µM), anti-ADAMTSL3 (250 ng/ml), anti-CADM1 (625 ng/ml), anti-GULP1 (625 ng/ml), anti-MDGA2 (62.5 ng/ml), anti-PILRα (1.25 µg/ml) or anti-PTPRK (1.25 µg/ml). Infection levels, quantified by luciferase activity, are expressed relative to untreated controls (100% infection). Each color-coded dot indicates an individual tissue with median and interquartile range indicated for each treatment with dotted lines. Statistical significance was assessed using a Kruskal–Wallis test for multiple comparisons (****P < 0.0001, ***P < 0.001, **P < 0.01, *P < 0.05). (**C**) Plots showing Spearman correlation coefficients and associated significant P values comparing inhibitor effects across lung and kidney tissues.

Renal samples from cortical biopsies exhibited a distinct profile (***Figure 4B***). Inhibition of ACE2 and TMPRSS2 in this tissue was less robust and more variable compared to lung samples, with median values of infection reduction of 68% [39-82] and 60% [51-72] respectively (***Figure 4B***). Notably, inhibition of the JAK-STAT pathway with Ruxolitinib was more efficient than inhibiting canonical factors, significantly restricting viral entry with a median of 84% reduction ([61-93], ***Figure 4B***). Moreover, in contrast to lung derived cells, IL1R1 and ADAM17 inhibitors showed a variable and mild effect, with median values around 20-24% reduction, which were similar to other candidates with low impact in these samples such as anti-MDGA2, anti-GULP1, anti-CADM1 and anti-PTPRK (***Figure 4B***). Among the rest of the candidates, blocking ADAMTSL3 markedly reduced viral entry in some samples, but this effect was highly variable among samples, while PILRα had no effect (***Figure 4B***).

Additionally, we performed a correlation analysis of the capacity to limit viral entry among the drugs, when enough drug pairs were available (>4), to provide insights into potential associations between novel candidate molecules and canonical entry factors (***Figure 4C*)**. In lung derived cells, the blocking effect observed for anti-ADAMTSL3 positively correlated with the effect of blocking TMPRSS2 (p=0.034, ***Figure 4C***), while anti-ADAMTSL3 positively correlated with the effect of anti-PILRα (p=0.017, ***Figure 4C***). In renal samples, blocking ADAM17 or IL1R1 correlated with the effect of blocking TMPRSS2 (p=0.048 and p=0.003, respectively) and between themselves (p=0.017; ***Figure 4C***), regardless of the low benefit of blocking these molecules in this tissue compared to the lung one. Together, these results highlight that, multiple proteins, variably expressed among tissues and individuals, contribute to SARS-CoV-2 viral entry. We also highlight differences among the contribution of these candidates, including canonical receptors, in lung versus renal cortex samples, where inhibition of the JAK-STAT pathway showed the highest benefit among tested molecules.

### *In silico*-evaluation of the selected candidates

Lastly, we assessed the potential direct interaction between identified targets and the Spike protein, as additional data to support candidates as direct entry factors or modulators of other factors. In brief, we retrieved protein structures from the Protein Data Bank or modeled the atomistic structure with AlphaFold2 (***Figure 5A***). ADAMTSL3 was excluded for not having PDB structure and low-quality AlphaFold models. GULP1 was excluded because its globular domain resides in the cytoplasm, leaving only a short extracellularly accessible peptide, and JAK1 was also excluded as it is exclusively cytoplasmic (intracellular). For the rest of the candidates, protein-protein docking followed by Monte Carlo simulations lead to protein-protein poses with high interaction affinity, whose stability in solution was assessed with molecular dynamics (MDs) simulations. Across all spike-target pairs, we identified poses exhibiting strong interaction affinities with protein interaction energies, measured in PELE, below -110 kcal/mol (***Figure 5B***), encompassing both the open and close S1 or full S trimeric conformations. When we further analyzed the stability of such interaction using molecular dynamics, PILRα and PTPRK showed the highest number of interacting poses with S or S1 closed trimers, with several others demonstrating stable L-RMSD and I-RMSD values in MD simulations (≤15–20 Å) (***Figure 5C*** and ***Figure S9***), thus identified as potential spike-binders. ADAM17, CADM1, and MDGA2 were also identified as binders, although exhibiting less complementary stability compared to PILRα and PTPRK. ADAM17 bound to both open and closed S1 subunit and the closed trimer, while CADM1 and MDGA2 bound to the closed S1 trimer (***Figure S9***). Notably, certain ADAM17-Spike and CADM1-Spike complexes displayed lower overall stability due to the high mobility of distal interacting regions, despite maintaining stable interfaces (i.e., high L-RMSDs and low I-RMSDs) (***Figure 5C*** and ***Figure S9***). In contrast, IL1R1 yielded fewer high affinity poses, and the only observed pose corresponded to a non-binding configuration (***Figure 5D***). Although some candidates could not be evaluated, *in silico* predictions identified PILRα and PTPRK as potential interactors with the spike protein, followed by ADAM17, CADM1, and MDGA2, which were predicted to have fewer stable interactions.

**Figure 5.**
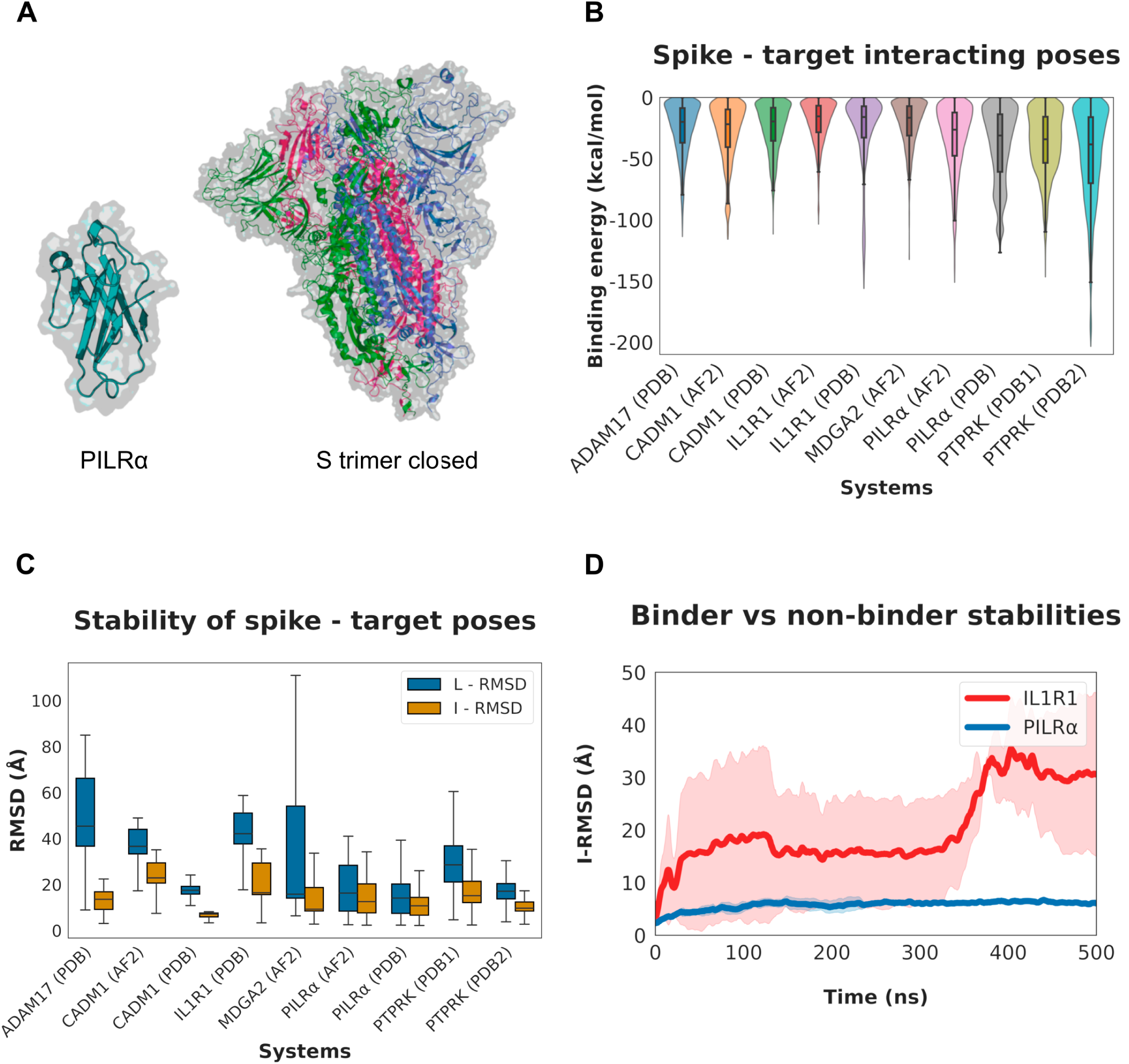
Computational modeling of spike–target interactions. **(A)** Examples of a protein monomer and spike conformation used in this study: PILRα and the spike protein in the closed conformation (colors correspond to each protein subunit). **(B)** Distribution of binding energies obtained from PELE simulations for each spike–target pair, across all spike conformations. **(C)** Distribution of RMSD values from molecular dynamics simulations of the top-ranked (strongest-affinity) poses identified in the previous stage. For each spike–target pair, RMSD data are pooled across all simulated target-spike conformation poses (individual data and concrete spike conformation in Supplementary Materials). **(D)** Stabilities of a binding (PILRα) and non-binding (IL1R1) target receptors, described using I-RMSDs from molecular dynamics simulations. Each curve represents the average I-RMSD metric obtained from the three independent replicas after smoothing, and the shaded area indicates the standard deviation among replicas.

## DISCUSSION

In this study, we provide a tissue-resolved characterization of SARS-CoV-2 entry mechanisms that extend beyond the canonical ACE2-TMPRSS2 axis. By integrating single-cell transcriptomics with functional pseudovirus assays in freshly isolated human epithelial cells, we show that viral entry is governed by a broader and context-dependent repertoire of membrane receptors than previously appreciated. The identification of regulators of inflammation, such as IL1R1, that highly impact viral entry in the lung parenchyma, and JAK1/2 in renal samples, supports a model in which SARS-CoV-2 exploits distinct molecular landscapes across organ systems together with the antiviral response. Furthermore, additional receptors heterogeneously expressed across samples and cell types, yet supported by genomics and computational analyses, may also contribute to mediate infection in certain individuals. Taken together, these findings help reconcile clinical observations of heterogeneous organ involvement and variable susceptibility across tissues and individuals.

Protein and gene signatures identified the most susceptible epithelial cells to viral entry across tissues, including ITGA6⁺ basal and suprabasal cells in nasal mucosa^41,42^, HLA-DR⁺/ITGA6⁺ AT2 cells in lung parenchyma^7^, and CD10⁺/CD13⁺ PTEC in kidney^4,39^. However additional cell types in the respiratory tract, including ciliated from nasal brushes and basal, multicilliated or even fibroblast from lung samples are also permissive to viral entry, although less efficiently. While these cell types have been established as the principal SARS-CoV-2–susceptible populations in their respective tissue^4,8,10–12,14,39^, the absence of enrichment for ciliated cells among permissive cells in our nasal samples^12^ likely reflects sampling constraints ^45,46^.

In this study, together with well-established canonical molecules that mediate SARS-CoV-2 viral entry, we confirm ADAM17 ^20,21,57^ as a significant contributor to viral uptake in cell targets derived from lung parenchyma, mostly identified as AT2. ADAM17 has been shown to cleave SARS-CoV-2 spike *in vitro*, promoting SARS-CoV-2 cell entry in a TMPRSS2-independent manner, likely modulating the lysosomal entry pathway ^20^. This same study showed that inhibiting this pathway markedly reduced infection of primary human bronchial epithelial cells ^20^, which now we extend to lung-derived AT2 cells. An additional proposed mechanism for ADAM17 enhanced infectivity could be through increasing ACE shedding, as reported ^21^. The lack of correlation between ADAM17 and ACE2 in the lung, as well as the positive correlation detected for the effect of blocking ADAM17 and TMPRSS2 in kidney, does not directly support any of these mechanisms. We also uncover a previously unappreciated role for IL1R1 in lung-derived primary cells. IL1R1 signaling can enhance ADAM17 activity and vice versa^58^, which is supported by our data showing a strong correlation on the effect of blocking these two molecules, yet only detected in renal samples. Further, inflammatory cytokines released in severe COVID-19, including IL-1β, have been shown to upregulate ACE2 expression, potentially establishing a positive-feedback loop for viral replication^3,9^; while using IL1R antagonists reduced the requirement of mechanical ventilation and decreased mortality in patients with COVID-19^59^. Here, IL1R1 blockade markedly reduced viral entry in lung, supporting early therapeutic targeting in high-risk groups.

In renal cortex, pseudovirus positive epithelial clusters revealed CADM1, PTPRK, and IL1R1 as candidate factors, consistent with prior observations of IL1R1 upregulation in renal epithelial cell lines after SARS-CoV-2 exposure, which contributes to cell injury without viral replication^60^. However, blocking IL1R1 or ADAM17 produced modest, variable effects that paralleled reductions in TMPRSS2-dependent inhibition, particularly weaker in kidney than lung, indicating tissue-specific mechanisms linking these molecules to viral entry. Notably, inhibition of the JAK-STAT pathway with Ruxolitinib was the most effective strategy to restrict viral entry in renal epithelial cells. Although JAK inhibition dampens IFN responses^35,61^, it also limits ACE2 and TMPRSS2 upregulation^9,62^, aligning with our functional data. However, we did not detect any correlation between the effect of Ruxolitinib and the effect of Camostat or anti-ACE2. Contrastingly, JAK1/2 blockade had no effect in lung-derived cells. Of note, these two tissues also differed in their dependency on ACE2 and TMPRSS2 canonical mechanisms. In kidney, SARS-CoV-2 can exploit receptor-mediated endocytosis through interaction between spike and secreted ACE2, alone or combined with vasopressin^21^. Indeed, activation of JAK during infection may further enhance ACE2 shedding via ADAM17 and ADAM10^18–21^, favoring this alternative entry route, which appears more effective in kidney. These tissue specific mechanisms may be particularly relevant for individuals with pre-existing renal vulnerability, who are at increased risk of severe or prolonged COVID-19.

Regarding remaining candidate receptors emerging from tissue specific analyses displayed variable but informative effects. CADM1 blockade reduced infection in selected renal and lung samples, whereas PTPRK inhibition produced modest yet consistent reductions in kidney. Using murine models and computational modeling, CADM1 was previously shown to bind to S1 spike protein^63^, consistent with our data, although with much less affinity than ACE2. Instead, PTPRK showed several stable interactions as predicted by our computational analyses. No study has suggested a role for this protein tyrosine phosphatase receptor before, although this gene, *PTPRK,* was one of the 14 genes common to SARS-CoV disease and cancer, cardiac, immunological and respiratory diseases^64^. Additional candidates, including MDGA2, GULP1, ADAMTSL3, and PILRα, showed divergent tissue-dependent roles. No literature has previously underscored a potential role for any of these proteins modulating or participating in any way in viral entry, except for -, which was described as a co-receptor for HSV-1^56^ and our *in silico* analyses suggested stability of various interacting poses with the spike. Targeting PILRα, a candidate obtained from basal respiratory cells, showed a clear benefit in most samples from lung and none from kidney. In fact, in lung samples, the inhibitory effect of PILRα positively correlated with the effect of blocking ADAMTSL3, which in turn correlated with inhibiting TMPRSS2. Mechanistically, the inhibitory immunoreceptor PILRα seems interesting because it modulates innate immune signaling and binds sialylated ligands, which have been shown to be involved in coronaviruses entry, including SARS-CoV-2^65^.

Overall, our findings reveal organ specific receptor usage and inflammatory interactions that shape SARS-CoV-2 entry in human epithelia. Targeting accessory receptors or inflammatory pathways in lungs may complement ACE2 or protease directed approaches, while modulation of kidney associated factors, particularly JAK1/2, may protect individuals at increased risk of renal involvement. This work refines the mechanistic landscape of SARS-CoV-2 dissemination and supports the development of tissue tailored antiviral interventions.

### Limitations of the study

Among other limitations already mentioned along the manuscript, we used the original D614G strain of SARS-CoV-2 to study viral entry factors in multiple tissues, and not other variants like omicron or delta. The choice of this strain was based on relatively broad tissue tropism, including lung, gastrointestinal tract and kidney, and on preserved entry pathways, since this variant relies more heavily on ACE2/TMPRSS2 host factors^66–68^. Moreover, the use of a pseudovirus instead of a replication competent virus may not fully reproduce the complexity of viral entry. The intricacy of the experimental setup, coupled with the limited availability of human samples, precluded comparisons across multiple variants, restricted simultaneous within-sample testing of all compounds, and restricted determination of effective concentrations (EC₅₀). In addition, donor variability or slight differences of the specific tissue region sampled may have biased target cell proportions, affecting results consistency. However, compared to other models, results from blocking these targets may be closer to reality.

## ACKNOWLEDGMENTS

We would like to thank all the volunteers and patients who participated in the study and their providers. The authors thank Joan Punyet and Norberto Núñez, from the Flow Cytometry at VHIR and Marco Antonio Fernández from the one at Institut de Recerca Germans Trias i Pujol, for excellent technical assistance. Funding: This work was primarily supported by Fundació La Marató TV3 (2021121 FMTV3). This work was additionally supported with grants from MCIN/AEI/10.13039/501100011033 and the European Union «Next Generation EU»/PRTR (CNS2022-135549 and PID2023-149204OB-I00). C.M-P. was supported by a Ph.D. fellowship from the Agència Gestió Ajuts Universitaris i de Recerca (AGAUR/FI/2023). I.B. was supported by the predoctoral fellowship AGAUR-FI (2025 FI-1 00783) Joan Oró, funded by the Department of Research and Universities of the Government of Catalonia, with co-funding from the European Social Fund Plus. The funders had no role in the study design, data collection and analysis, the decision to publish, or preparation of the manuscript.

## AUTHOR CONTRIBUTIONS

M.V.R., C.M-P., A.B-M. and I.M. conducted infection assays. C.M-P., A.B-M. and DKJ.P. sorted cells from tissue models. MA.M. Performed single cell gene expression analyses which was supervised by H.H. I.B. and V.M. performed computational analyses which was supervised by V.G. BP, J.R., ME.S., S.L., E.E-B., E.T., V.F., A.C., selected study subjects and provided human samples which was supervised by J.B. M.V.R., A.G. and J.G-E. made pseudoviruses, which was supervised by MJ.B. M.G. conceptualized and supervised the project. J.B., H.H. and M.G. acquired funding. MJ.B., V.G. and H.H. contributed to the design and interpretation of the experiments. M.V.R. and M.G. wrote initial text drafts. All authors reviewed and edited the manuscript.

## SUPPLEMENTAL INFORMATION

Document S1. Figures S1–S9 and Tables S1-S4.

## STAR★METHODS

### KEY RESOURCES TABLE

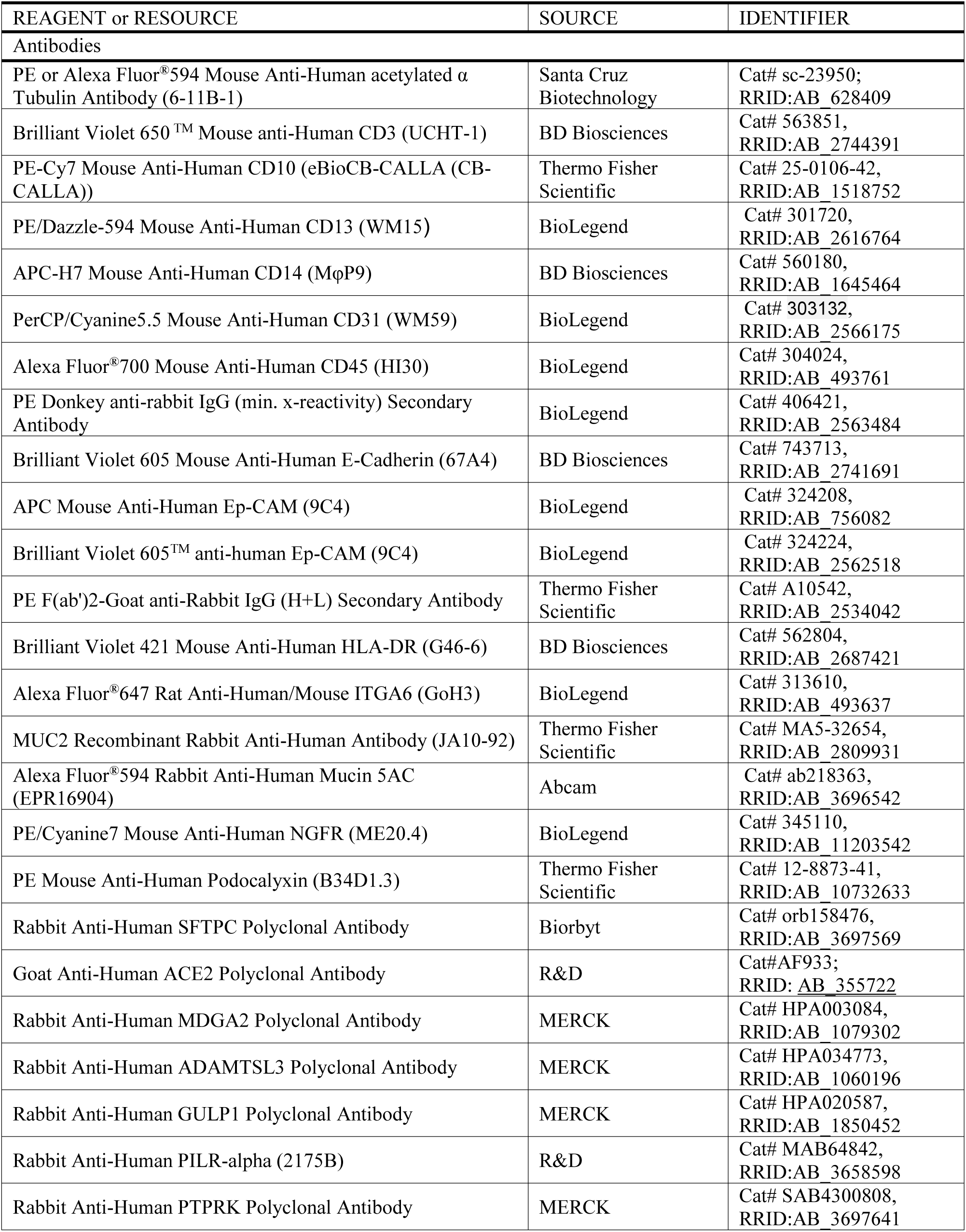
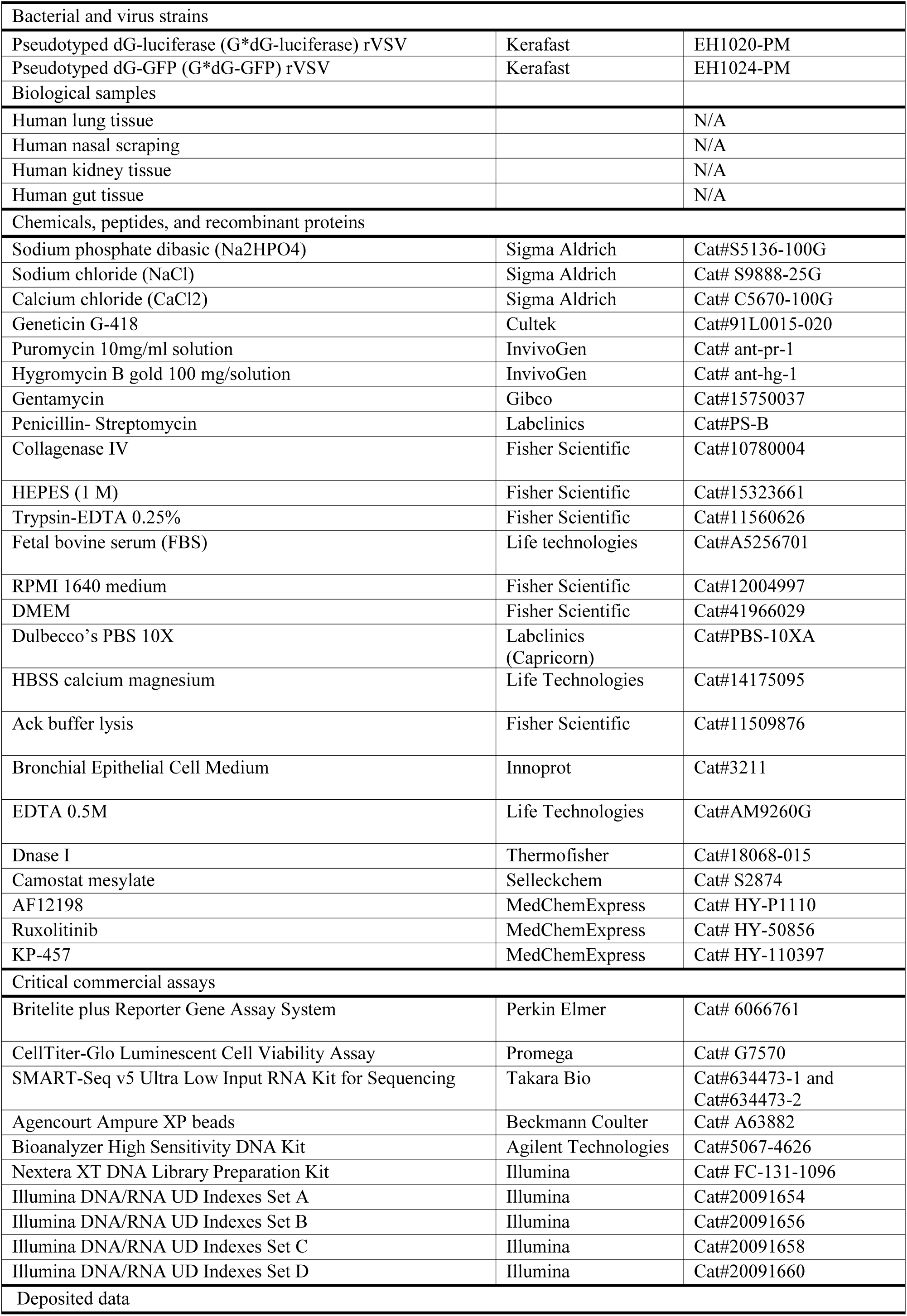
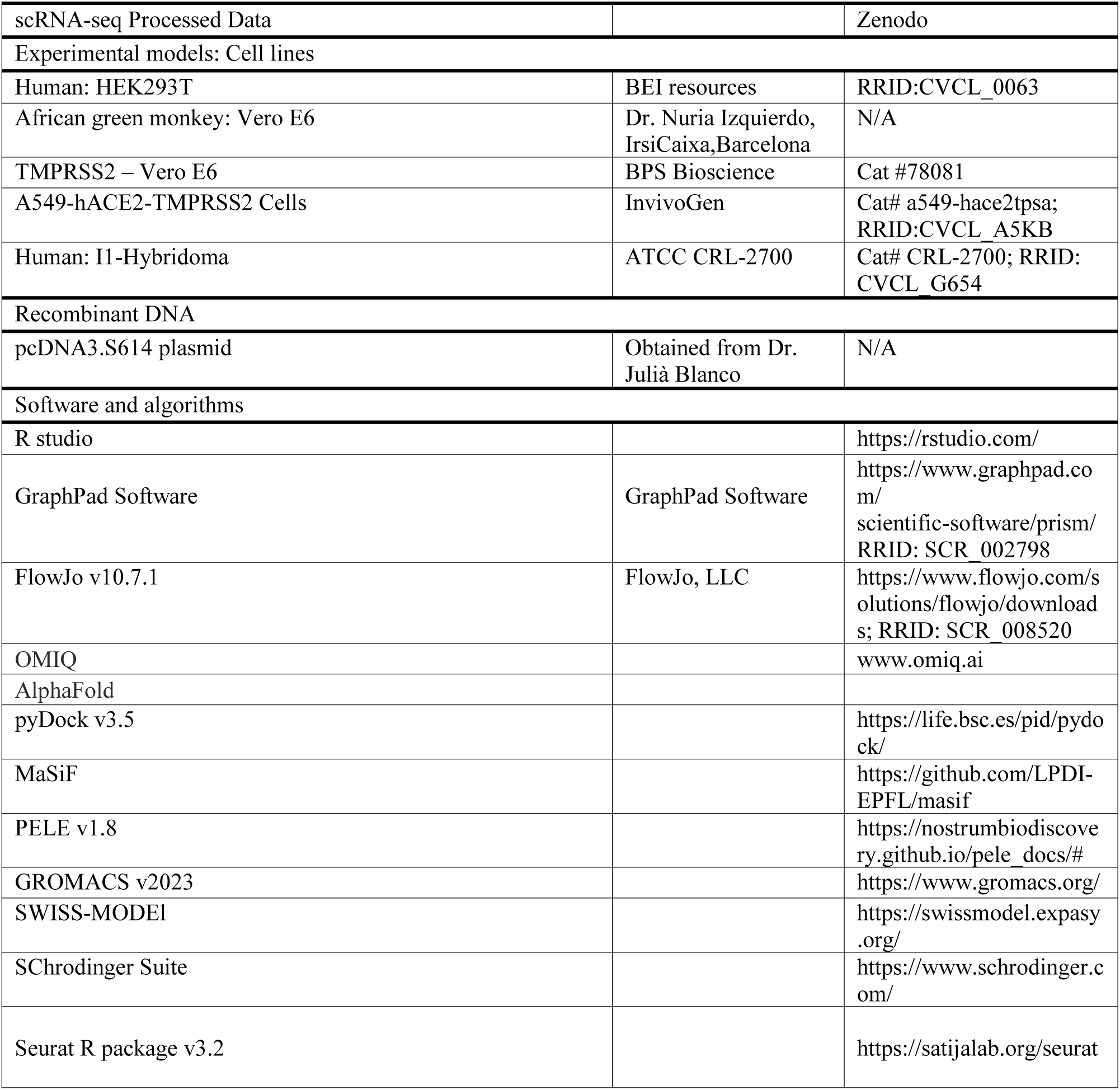

#### Experimental model and subject details

All cell lines were maintained at 37 °C under 5% CO_2_. HEK293T (BEI resources), Vero E6 (kindly provided by Dr. Nuria Izquierdo, IrsiCaixa Institute, Barcelona) and Vero E6-TMPRSS2 (BPS Biosciences) were cultured in Dulbecco’s Modified Eagle Medium (DMEM) supplemented with 10% fetal bovine serum (FBS; Gibco) 100 U/ml penicillin, and 100 μg/ml streptomycin (Capricorn Scientific). The A549-T hACE2/TMPRSS2 cell line (InvivoGen) was maintained under the same conditions with the addition of 150 mg/ml Hygromycin (InvivoGen) and 0.5 ug/ml of Puromycin (InvivoGen).

#### Study participants

The study protocol was approved by the Hospital Universitari Vall d’Hebron Institutional Review Board (Comitè d’Ética d’Investigació Clínica; IRB number PR(AG)45/2021) Barcelona, Spain. To ensure adherence to ethical standards, all participants provided written informed consent prior to sample collection. Healthy human nasal epithelial samples, lung, kidney, and gut tissues were obtained from adult male and female donors and patients with no COVID-19 symptomatology through established collaborations with different clinical departments from Hospital Universitari Vall d’Hebron (HUVH). Patient clinical characteristics are detailed in Table S1.

## Method details

### Human Tissue Sample processing

Fresh lung and kidney tissues were preserved in RPMI-1640 medium supplemented with antibiotics, dissected into 2 mm-thick blocks, and digested with 5 mg/mL collagenase IV (Gibco) and 100 μg/mL DNase I (Roche, Basel, Switzerland) for 30 min at 37°C with agitation (400 rpm). Colon samples was incubated overnight in RPMI-1640 medium with antibiotics and antifungals prior to dissection and digestion with 2.5 mg/mL collagenase IV and 100 μg/mL DNase I for 30 min. Digested tissues were mechanically disrupted using a pestle, then filtered through 70 μm cell strainer (Labclinics) and washed twice with PBS. For lung and gut, an additional incubation with 100 μg/mL DNase I in 5 mL HBSS+/+ (ThermoFisher) was performed, followed by titration through a 40 μm pore size cell strainer (Labclinics). Cell suspensions were counted using a LUNA-FL™ Dual Fluorescence Cell Counter (Logos Biosystems). Nasal epithelial cells were obtained with an ASI Rhino-Pro® nasal curette (Arlington, IL, USA) into Bronchial Epithelial Cell Medium (BEpiCM) (Innoprot) washed with PBS with 5 mM EDTA, incubated on a shaker (15 min. 30 rpm, 4°C) centrifuged, filtered and counted.

### Pseudovirus production

Pseudotyped viral stocks of VSV*ΔG(GFP)-S(D614G) and VSV*ΔG(Luc)-S(D614G) were generated following the protocol described by Whitt^69^, with further modifications implemented by our laboratory^7^. Briefly, 293T cells were transfected with 3μg of the plasmid encoding the SARS-CoV-2 spike (pcDNA3.S614). Next day, cells were infected with a VSV-ΔG-GFP or VSV-ΔG-Luc virus (Multiplicity of Infection, MOI = 1) (generated from a lentiviral backbone plasmid that uses a VSV promoter to express luciferase) for 2h and gently washed with PBS. Cells were incubated overnight in D10 supplemented with 10% of I1 hybridoma (anti-VSV-G) supernatant (ATCC CRL-2700) to neutralize contaminating VSV*ΔG(Luc/GFP)-G particles. Next day, the resulting viral particles were collected and titrated in VeroE6/TMPRSS2 (BPS Biosciences) cells using a luminescence-based enzyme assay (Britelite Plus Kit; PerkinElmer), or flow cytometry for GFP viruses.

### SARS-CoV-2 Pseudovirus -GFP Infection

Cell suspensions from nasal, kidney and gut tissues were seeded at 3.5x10^5^ cells/well in 96-well conical-bottom plates; lung cells were seeded at 7x10^5^ cells/well. Cells were then infected with VSV*ΔG(GFP)-S virus, and centrifuged at 1,200 g, at 37 °C for 30 min (nasal and kidney) or for 2 h (lung and gut). Following infection, cells were resuspended in fresh medium, transferred to a 96-well flat-bottom plates, and cultured overnight at 37°C in a 5% CO2 incubator.

### Spectral Flow cytometry Phenotyping and Sorting

The following day, kidney suspensions underwent erythrocyte lysis with ACK Lysing Buffer (250µl; Gibco). Cellular suspensions were later stained with Live/Dead® Aqua (Invitrogen), followed by surface staining with the antibodies required for phenotype identification. Common markers across all tissues included anti-CD31 (PerCP-Cy5.5, BioLegend), anti-CD14 (APC-H7, BD Biosciences), anti-CD45 (AF700, BioLegend), anti-EpCAM (BV605 and APC, BioLegend), anti-CD3 (BV650, BD Biosciences) and anti-HLA-DR (BV421, BD Biosciences). Infected cells were detected using GFP expression. For nasal samples we additionally included anti-NGFR (PE-Cy7, BioLegend), anti-MUC5AC (AF594, Abcam), anti- α-tubulin (PE, Santa Cruz Biotechnology) and anti-ITGA6 (AF647, BioLegend). For lung, anti-NGFR, anti-α-tubulin, anti-ITGA6 and primary anti-SPC (unconjugated, Biorbyt) with secondary donkey anti-rabbit IgG (PE, BioLegend). For kidney, we included anti-CD10 (PE-Cy7, ThermoFisher), anti-CD13 (PE/Dazzle594, BioLegend), anti-podocalyxin (PE, ThermoFisher), and anti-E-cadherin (BV605, BD Biosciences); and for colon, anti-NGFR, anti-E-cadherin and primary anti-MUC2 (unconjugated, ThermoFisher) with secondary goat anti-rabbit IgG (PE, ThermoFisher). After staining, cells were fixed in PBS 2% paraformaldehyde (PFA) and acquired in an Aurora spectral Cell Sorter (Cytek). Sorting was performed on the Cytek Aurora Cell Sorter directly in to SMART-seq2 plates previously conserved at -20°C. Phenotypic data analysis was conducted using FlowJo v10.7.1 (Tree Star) and Omiq software (Dotmatics, www.omiq.ai). Sorted plates were maintained at 4°C during the sorting and immediately after, covered with a lid, spined at 2000 rpm at 4°C and snap freezed on dry ice. Sorted plates were stored at 80°C until shipment. Later, plates were sent in dry ice to the Single Cell Genomics group at CNAG-CRG, Parc Científic de Barcelona.

### Single cell data generation and analysis

Optimized Smart-seq2 protocol with Takara SMART-seq® Single Cell Kit reagents for improved sensitivity and gene identification. Briefly, full-length single-cell RNA-seq libraries were prepared using the SMART-Seq v5 Ultra Low Input RNA Kit for Sequencing (Takara Bio). All reactions were downscaled to one quarter of the original protocol (after implementation of low-volume robotics) and performed following thermal cycling manufacturer’s conditions. Freshly harvested single cells were sorted into 96-well plates containing 2.5 µl of the Reaction buffer (1x Lysis Buffer, RNase Inhibitor 1U/µl). Reverse transcription was performed using 2.5 µl of the RT MasterMix (SMART-Seq v5 Ultra Low Input RNA Kit for Sequencing, Takara Bio). cDNA was amplified using 8 µl of the PCR MasterMix (SMART-Seq v5 Ultra Low Input RNA Kit for Sequencing, Takara Bio) with 25 cycles of amplification. Products of each well of the 96-well plate were pooled and purified twice with Agencourt Ampure XP beads (Beckmann Coulter). Final libraries were quantified and checked for fragment size distribution using a Bioanalyzer High Sensitivity DNA Kit (Agilent Technologies). Pooled sequencing of Nextera libraries was carried out using a NovaSeq6000 (Illumina) to an average sequencing depth of 0.5 million reads per cell. Sequencing was carried out as paired-end (PE150) reads on a NovaSeq6000 S4 flow cell with library indexes corresponding to cell barcodes.

### Preprocessing (Mapping and Quantification)

In the initial phase, we utilized the zUMIs^70^ single-cell RNA-sequencing pipeline to meticulously enhance the quality and reliability of our raw sequencing data. We initiated this process with quality control assessments aimed at eliminating low-quality reads and excluding any potential contaminants. Subsequently, we aligned the preprocessed reads to the GRCh38 reference genome, quantifying gene expression counts for each individual cell. These expression counts were subsequently subjected to normalization through log-transformation, specifically using the NormalizeData() function from Seurat^71^ with a size factor set to 10,000. This approach effectively accounted for any technical biases, ensuring that our data was suitably prepared for subsequent analyses.

### Feature selection, dimensionality reduction

To identify and annotate cell types, we initiated the process by employing the FindVariableFeatures() function to isolate the top 3,000 most highly variable genes, which effectively capture the major axes of biological variability within our dataset. Following this selection, we further standardized the data by applying a Z-score transformation using the ScaleData() function in Seurat, centering it at 0. We retained the top 20 principal components derived from principal component analysis (PCA) for use in subsequent analytical steps.

### Clustering and annotation

To facilitate the clustering process, we initiated it by constructing a K-nearest neighbors (KNN) graph through the utilization of the FindNeighbors() function. Subsequently, we employed the Leiden community detection algorithm to partition this graph. To optimize the clustering results, we selected a slide-specific resolution 0.2 value when executing the FindClusters() function in Seurat. Using unsupervised clustering methods, we effectively organized cells based on their distinctive gene expression profiles, thereby enabling the identification of discrete cell populations. Our approach to characterizing epithelial cell types revolved around the assessment of marker gene expression, including genes such as EPCAM, KRT19, and KRT18. Furthermore, to enhance the precision of our subtyping efforts, we performed a differential expression (DE) analysis specifically between the identified epithelial cell clusters. This DE analysis allowed us to further refine the categorization of epithelial cell subtypes based on the unique and discernible expression patterns of specific marker genes.

### Differential Expression (DE) Analysis Between GFP^+^ and GFP^-^

DE genes between GFP^+^ and GFP^-^ cells for each tissue and each cluster were identified by utilizing the FindMarkers function within the Seurat framework. To gain a deeper understanding of the biological significance of these differentially expressed genes within each tissue, we carried out functional enrichment analyses. Selection of gene candidates was based on established thresholds: average log₂ fold change (FC above >0.25) and adjusted p value (p_adj;_ <0.05). Differential expression results were visualized using the Enhanced Volcano R package (version 1.28.2).

### Assessment of antiviral activity of candidate inhibitors

Candidate inhibitors were tested in triplicate in human lung tissue (HLT) and kidney tissue (HKT). HLT cells were seeded at 3x10^5^ cells/well and incubated for 1h at 37°C prior to infection with VSV*ΔG(Luc)-S (MOI=1) as previously described by our lab^7^. Infection was facilitated by spinoculation at 1,200g for 2 h at 37°C. After infection, cells were resuspended in RPMI + 20% FBS and transferred to 96-well flat-bottom plates.

HKT cells were processed under the same conditions, except that spinoculation was performed for 1h. Each assay plate included the following controls: background (no cells), mock-infected cells and infected cells but untreated. At 20 h post-infection cells were incubated with Britelite Plus reagent (PerkinElmer) and subsequently transferred to an opaque black plate. Luminescence was immediately recorded on a plate reader (LUMIstar Omega). Infection was considered if luciferase activity (RLUs; relative light units) was above 2.5 times the RLU in the mock-infected wells. Cytotoxicity was evaluated in parallel in some samples, using the CellTiter-Glo Luminescent kit (Promega), following the manufacturer’s instructions.

### In-silico analysis system preparation

Protein hits obtained from the RNA seq analyses were analyzed using computational simulations to explore the potential direct protein-protein interaction between the identified targets and the Spike protein. For every identified target, we downloaded their available protein structure from the PDB database (ADAM17: 8SNL, CADM1: 4H5S, IL1R1: 4GAF, PTPRK (1): 2C7S, PTPRK (2): 8A1F). Moreover, for every target we used AlphaFold2.3^72^ to predict their 3D structure, and retain those models with sufficient quality (based on plDDT and pTM scores) or presenting complementary conformations to the available PDB structure. In addition, we downloaded the 3D structure from the Spike protein trimer in its open (6VYB) and closed (6VXX) conformations. Notably, we recovered missing atoms/residues using ColabFold^73^ and Swiss-Model^74^ (for those systems with extreme large size). Right after, we run a short molecular dynamics simulation (500 ps; full details in Molecular dynamics section-) to energetically accommodate the protein structures and minimize small artifacts.

### Rigid protein-protein docking

Rigid-body protein–protein docking simulations were performed using PyDock v3.5^75^ with default parameters. For each spike–target protein pair, the target protein (receptor) was kept fixed, while the spike protein (ligand) was systematically rotated and translated to explore possible binding orientations. A total of 92,400 docking poses were generated for each pair of proteins using the FTDock sampling algorithm implemented in PyDock. Atop from the full-length spike trimers, we generated a complementary trimer structure restricting to include only the S1 domain. Such decision was made to increase the density of sampling in the RBD, which has been widely described to be implicated in multiple protein-protein interactions. We also decided to keep the full-length under the assumption to identify potential binding spots beyond the RBD.

### Protein-protein interface filtering and clustering

Then, a filtering strategy was applied to only keep a set of non-redundant poses with potential for protein-protein interactions. Firstly, interacting regions on spike and target proteins independently, were predicted using the Molecular Surface Interaction Fingerprinting (MaSIF)-site method ^76^ with default parameters. The resulting prediction maps, binarized at probability 0.7, were clustered using k-Means selecting the optimal number of clusters based on a silhouette analysis. Only docking poses with proximal interacting regions were kept: at least 30% of the spike residues were in contact with any residue of the target protein were retained, considering a contact to occur when two Cα atoms, one from each chain, were within 10.0 Å of each other. Since there could be more than one interacting region per protein, combinations of these were made independently and resulted in independent sets of protein-protein poses with partially overlapping interface residues.

The filtered protein–protein poses were then clustered to remove redundant conformations with similar interfaces while prioritizing those with favorable energy values (as lower energies reflect fewer steric clashes). The contact residues were computed for each docking pose, and a Jaccard index was calculated to quantify the similarity between contact sets. Poses were iteratively assigned to clusters by comparing only with the representative pose of each existing cluster, defined as the pose with the lowest energy within that cluster. A Jaccard index threshold of 0.3 was used to determine whether a pose belonged to an existing cluster or initiated a new one, ensuring that each cluster represented a distinct binding interface while maintaining energetically favorable interactions.

Finally, a set of approximately 50 - 200 poses resulted out of each set of protein - spike conformation docking sampling. These were prepared using the Protein Preparation Wizard tool from the Schrödinger suite v2025^77^ for subsequent analyses. Briefly, disulfide bonds between close cysteines were established, protonation states of titratable residues at pH 7.4 were calculated with PROPKA 3.0 and the hydrogen-bonding network was optimized based on those pKa values, and the structure was subjected to a restrained minimization, with a root mean square deviation (RMSD) maximum of 0.3 Å, using the all-atom OPLS2005 force field. Of note, we did not include the protein-protein interacting complexes from AI-based predictions using AlphaFold3^78^, given its low confidence scores (all target - S protein interfaces whether monomer or trimer had iPTM < 0.8).

### Refinement of protein-protein poses

Protein Energy Landscape Exploration^79^ (PELE, v1.8) simulations were performed for conformational exploration and refinement and binding affinity estimation of the spike proteins to the target. In short, PELE is a heuristic Monte Carlo (MC)-based software that samples microstates in protein-ligand or protein-protein complexes. Firstly, it applies small rotations and translations and incorporates protein flexibility through the normal modes of the anisotropic network model. Next, side chain rotamer prediction is performed at protein-protein interfaces to relieve steric clashes, and a truncated Newton minimization is applied using the OPLS2005 forcefield. The effect of water molecules is considered using the Variable Dielectric Generalized Born Non-Polar (VDGBNP) implicit solvent model. Finally, the sampled microstates are accepted or rejected based on the Metropolis criterion^80^ and total and binding energy (i.e. an estimation of interaction/binding affinity between the proteins) metrics are computed on the sampled poses, completing one Monte Carlo step.

For each spike–target protein pair, the spike protein was truncated to reduce system size, allowing the simulation to focus on the interface while improving computational efficiency. The truncation retained all residues whose Cα atoms were within 25.0 Å of any Cα atom of the target protein. Additionally, regions of up to 10 residues bridging gaps were included. Isolated residue chunks of up to 20.0 Å were removed, and short, distant loops were further excluded following visual inspection to ensure the truncated structure retained only relevant interaction regions.

We run PELE simulations on the resulting protein-protein complexes from previous steps in two stages: equilibration and production. In the equilibration stage, with the aim of correcting interface clashes derived from FTDock and obtain an adequate baseline pose (i.e. low system energy), we applied small perturbations (rotation factor of 0.01 rad and translation range of 0.02 Å) using 1–3 computing cores per pose for 50 MC steps based on the total number of poses: 3 cores for ≤100 poses, 2 cores for 101-200 poses, and 1 core for >200 poses. Sampled microstates per protein-protein pose were grouped and the pose with the lowest binding energy was selected as the equilibrated state of that pose. Then, the top 20 binding energy equilibrated poses were moved to production. In the production stage, higher perturbations (rotation factor of 0.01 or 0.04 rad and translation range of 0.25 or 0.5 Å) were used on the spike protein to model protein-protein interactions, and 4 cores per equilibrated pose were used for 200 MC steps. This produced a distribution of protein poses of the target receptor with the different spike conformations, where those with the lowest protein-protein interaction energies represent the strongest interacting poses.

### Molecular dynamics simulations

Molecular dynamics (MDs) simulations using Gromacs v2023.3^81^ and the force field CHARMM36^82^ were performed to assess the stability of the predicted poses with promising binding affinities estimations (i.e. binding energies). These were run on poses with strong binding affinities, defined as those with binding energy of -110 kcal/mol or less, allowing a maximum of 5 poses per spike-target pair. Briefly, we solvated a periodic octahedron box consisting of a TIP3P water model^83^ and Na+ and Cl– ions at 0.15 M. Then, we ran a steepest descent minimization and then 200 ps of constant number-volume-temperature (NVT) equilibration step of the water solvent, restraining the position of all atoms with a force constant of 1000 kJ/(mol·nm^2^). Afterwards, the system was heated to 300K by running ten 50 ps of constant number-pressure-temperature (NPT) MDs simulations using different protein residues constrains with force constants of 1000, 550, 300, 170, 90, 50, 30, 15, 10 and 5 kJ/(mol·nm^2^). Then, a production of 500 ns was run, using a 2 fs time step (at temperature 300 K, pressure 1 bar and particle-mesh Ewald for electrostatic interactions). For each protein-protein pose, 3 replicas of the previous MDs protocol were performed. The ligand RMSD (L-RMSD), defined as the RMSD of the target Cα atoms after aligning trajectory structures using the truncated spike protein, and the interface RMSD (I-RMSD), defined as the RMSD of protein–protein contact residues (residues with Cα atoms within 10.0 Å, including those included in gaps of one to three residues between contact residues), were computed to assess the stability of both the complex and the interface. Notably, the L-RMSD measures the deviation of the target structure from the initial PELE pose, with higher values indicating greater deviation; the I-RMSD reflects the stability of the interface relative to the initial PELE pose, where lower values correspond to a more stable interface.

### Statistical analysis

All graphing and statistical analyses were generated using GraphPad Prism version 10 (GraphPad Software, La Jolla, CA, USA) with statistical significance defined as p< 0.05. The statistical details and specific statistical test used for each dataset is mentioned in the respective figure legend.

---

**Figure S1.**
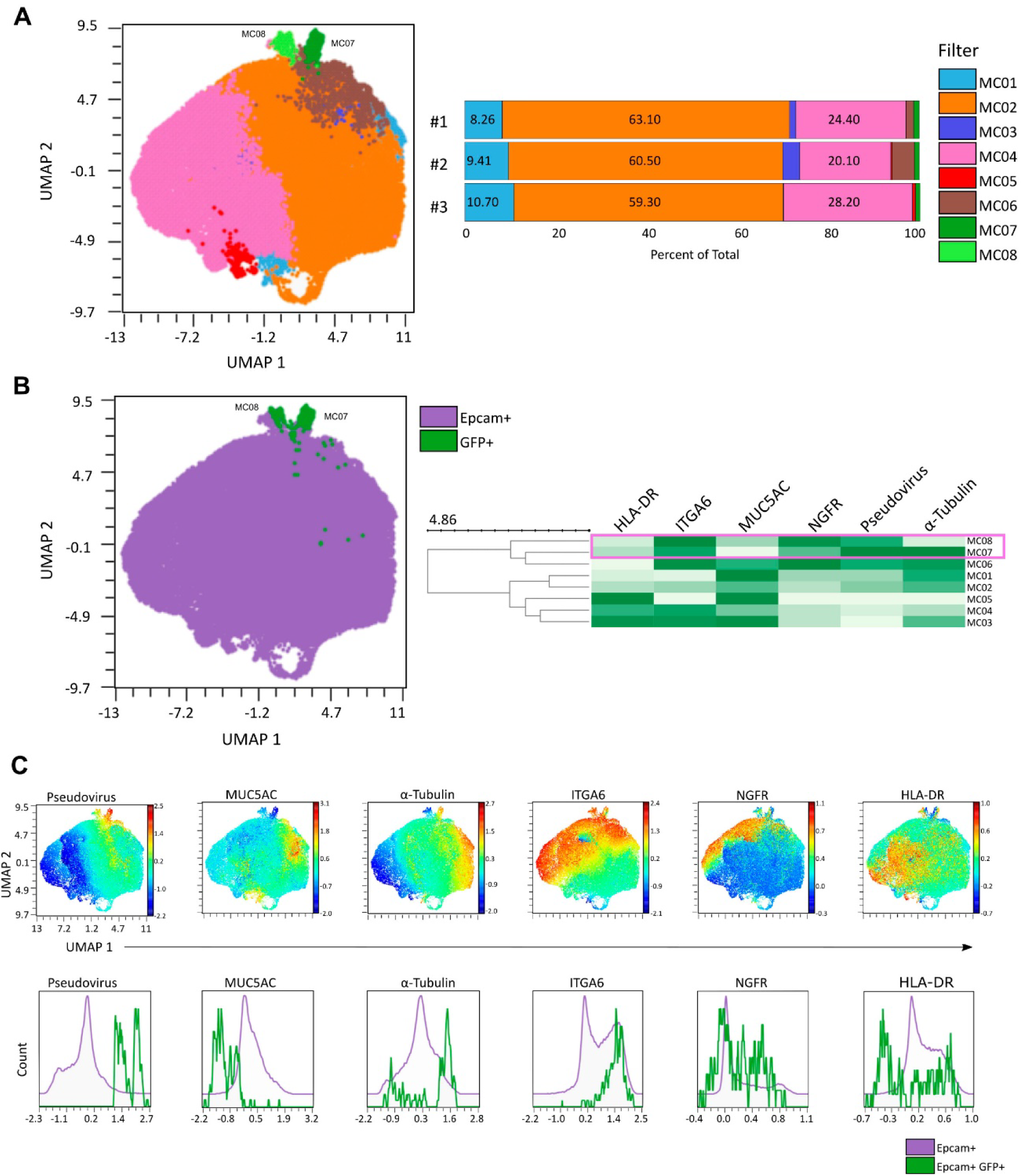
UMAP-based clustering and phenotypic characterization of infected nasal epithelial cells. **(A)** UMAP projection of high-dimension single-cell flow cytometry nasal data of EpCAM^+^ cells from *ex vivo* SARS-CoV-2 pseudovirus infected samples, depicting the differentiation of 8 clusters (MC) and relative abundance of each cluster across the three pools of nasal samples. **(B)** UMAP visualization highlighting the EpCAM^+^GFP^+^ infected population (green). Adjacent heatmap displays normalized mean fluorescence intensity of epithelial markers across the 8 identified clusters as indicated in **(A)**. **(C)** UMAP-based visualization of the spatial expression of six molecules (top) and density histograms comparing EpCAM⁺ and EpCAM⁺GFP⁺ populations (bottom).

**Figure S2.**
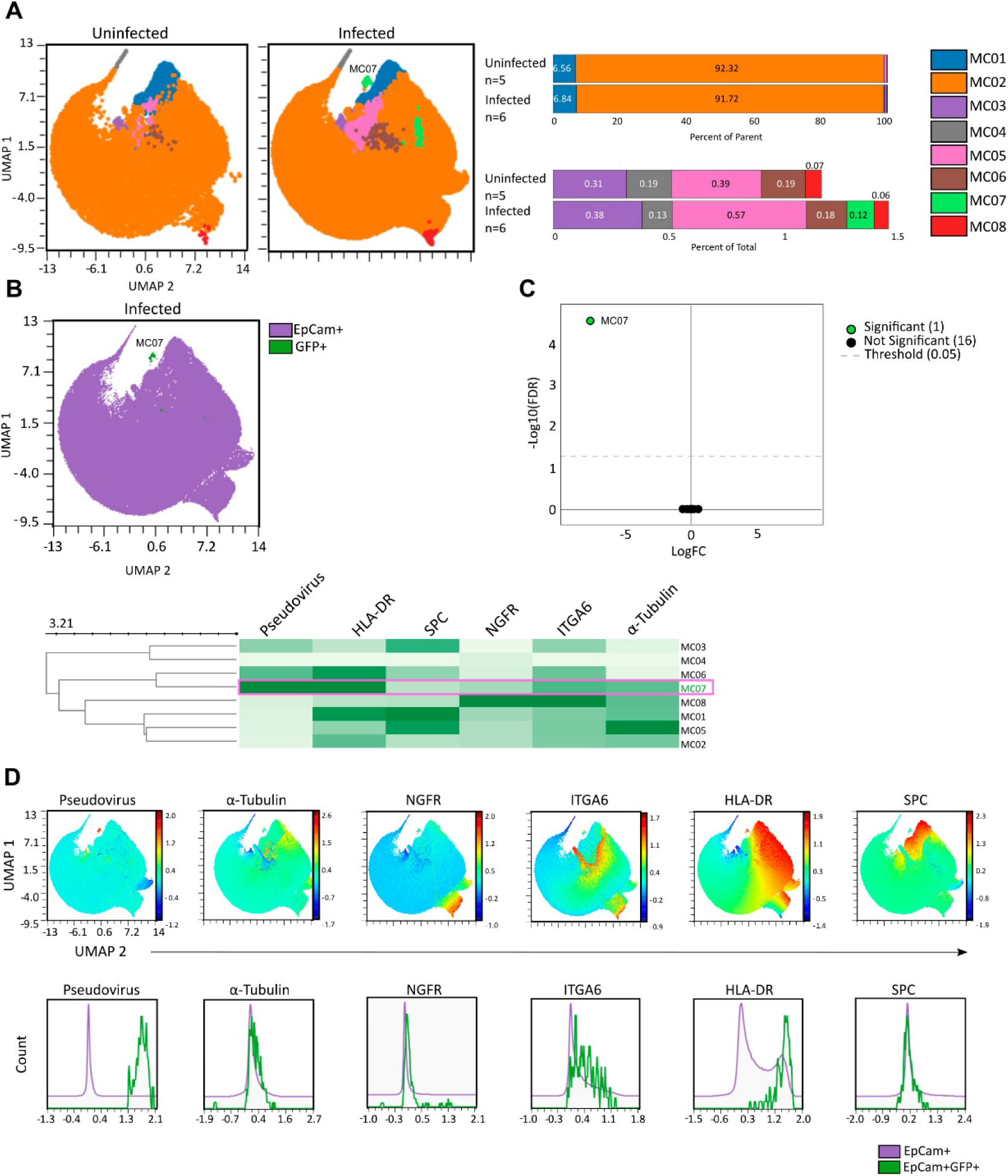
UMAP-based clustering and phenotypic characterization of infected lung-derived epithelial cells. **(A)** UMAP projection of high-dimension single-cell flow cytometry data of pulmonary EpCAM^+^ cells from uninfected or *ex vivo* SARS-CoV-2 pseudovirus infected samples, depicting the differentiation of 8 clusters (MC) and relative abundance of each cluster across uninfected versus infected samples (bars on top). Bars below show the percentage of less represented clusters (MC03-MC08). **(B)** UMAP visualization highlighting the EpCAM^+^GFP^+^ infected population (green). Adjacent heatmap (bottom) displays normalized mean fluorescence intensity of epithelial markers across the 8 identified clusters as indicated in **(A)** highlighting the cluster representing GFP^+^ cells (MC07). **(C)** Volcano plot displaying the differential cluster abundance comparing uninfected and infected samples, the green dot corresponds to MC07. **(D)** UMAP-based visualization of the spatial expression of six molecules (top) and density histograms comparing EpCAM⁺ and EpCAM⁺GFP⁺ populations (bottom).

**Figure S3.**
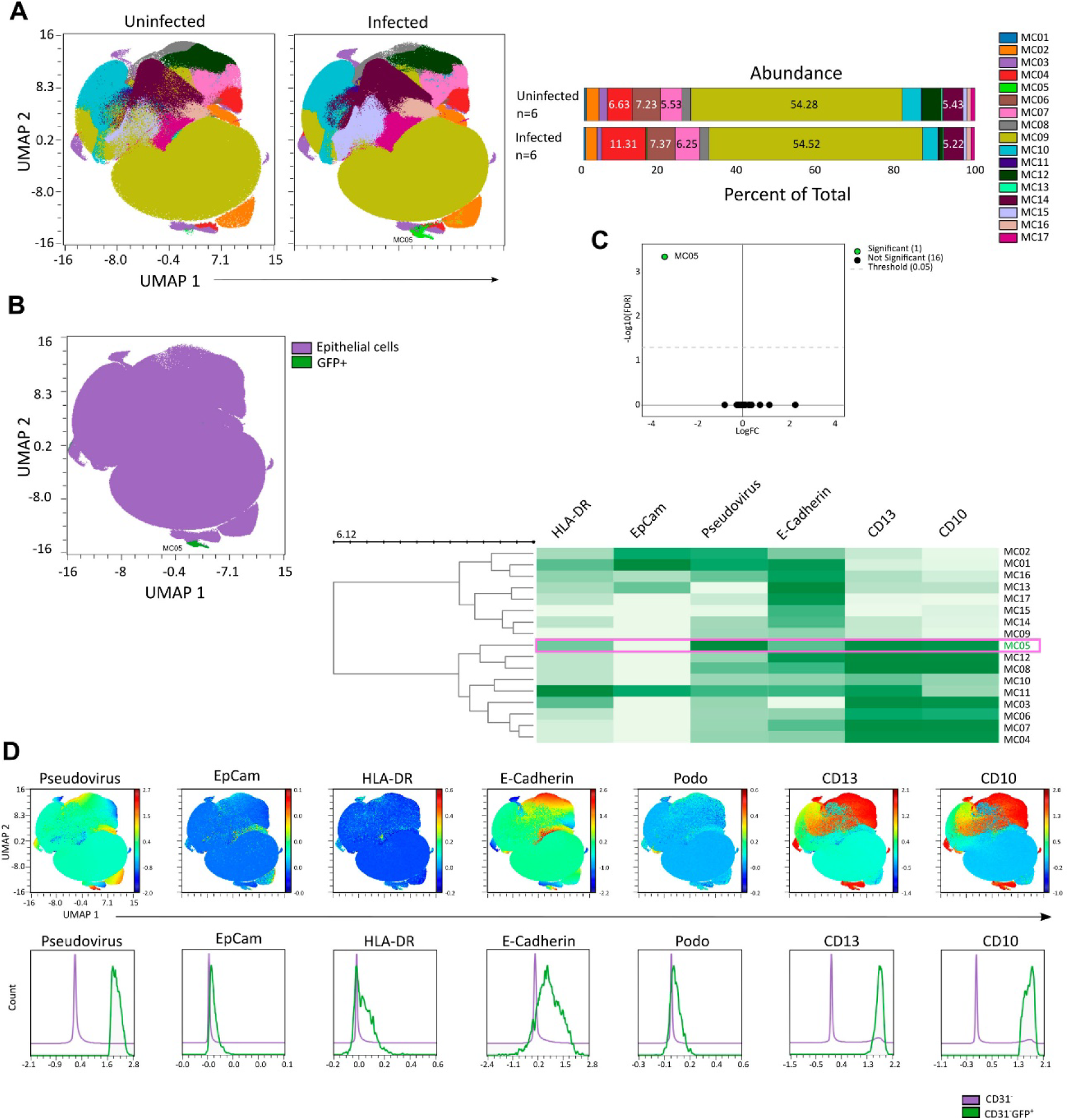
UMAP-based clustering and phenotypic characterization of infected renal cells. **(A)** UMAP projection of high-dimension single-cell flow cytometry data of renal CD31^-^ cells from uninfected or *ex vivo* SARS-CoV-2 pseudovirus infected samples, depicting the differentiation of 17 clusters (MC) and relative abundance of each cluster across uninfected versus infected samples. **(B)** UMAP visualization highlighting the CD31^-^GFP^+^ infected population (green). Adjacent heatmap displays normalized mean fluorescence intensity of epithelial markers across the 17 identified clusters as indicated in (**A**) highlighting the cluster representing GFP^+^ cells (MC05). **(C)** Volcano plot displaying the differential cluster abundance comparing uninfected and infected samples, the green dot corresponds to MC05. **(D)** UMAP-based visualization of the spatial expression of six molecules (top) and density histograms comparing CD31^-^ and CD31^-^GFP^+^ populations (bottom).

**Figure S4.**
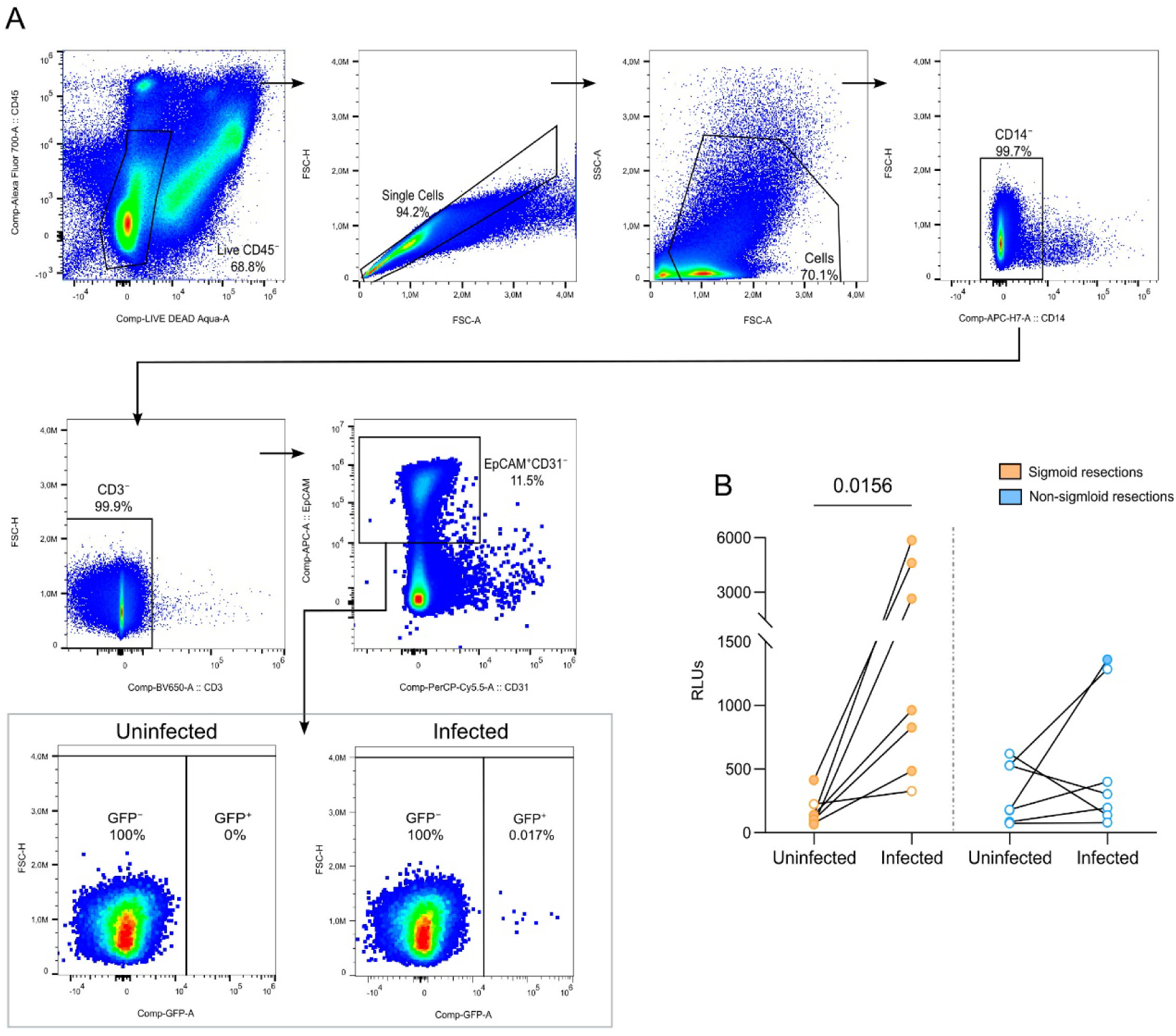
SARS-CoV-2 pseudoviral infection of human colon. Representative flow-cytometry plots illustrating the gating strategy used to assess ex vivo SARS-CoV-2 pseudoviral infection in human colon tissue, including detection of GFP⁺ cells in paired uninfected and infected samples. **(B)** Paired relative light unit (RLU) measurements compare uninfected and infected conditions in sigmoid (left, orange) and non-sigmoid (right, blue) colon resections. Each pair of dots represents an individual sample. Infection was defined as values exceeding 2.5-fold above the mean uninfected background. Infected samples were significantly enriched in sigmoid resections (bold dots), whereas non-sigmoid samples showed a higher proportion of non-infected cases (open dots).

**Figure S5.**
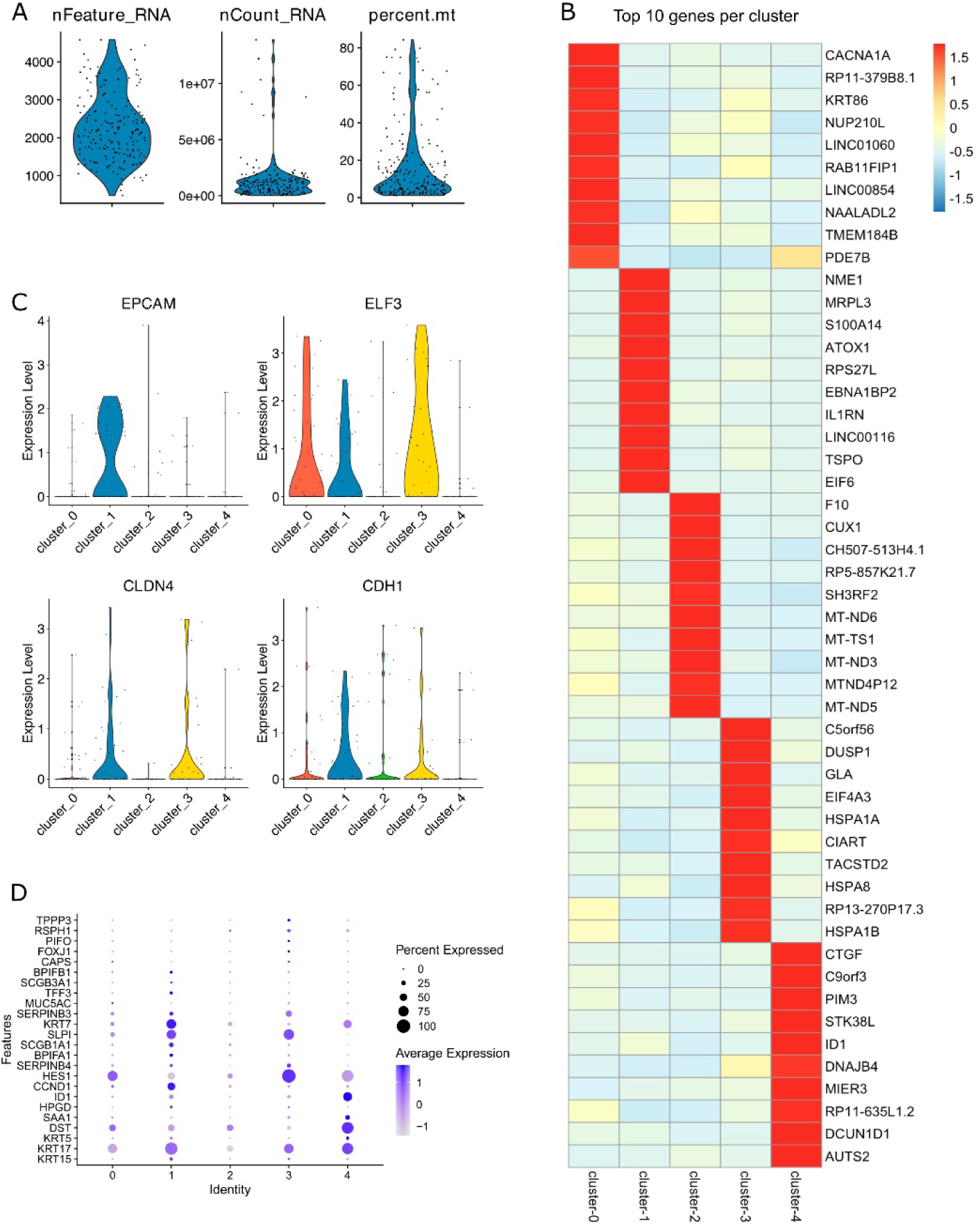
Single-cell RNA-seq quality control, clustering, and marker gene expression in nasasamples. **(A)** Violin plots showing distributions of standard single-cell RNA-seq quality control metrics across all cells, including the number of detected genes (nFeature_RNA), total UMI counts (nCount_RNA), and the percentage of mitochondrial transcripts (percent.mt). (**B**) Heatmap of the top 10 differentially expressed genes for each identified cluster. Genes are ranked by average log-normalized expression within each cluster relative to others. Color scale indicates scaled expression levels. (**C**) Violin plots depicting normalized expression levels of selected epithelial marker genes (*EPCAM, ELF3, CLDN4*, and *CDH1*) across the indicated clusters, highlighting cluster-specific expression patterns. (**D**) Dot plot summarizing the expression of representative marker genes across clusters, based on Ahn, J.H., et al. *(2021. J. Clin. Invest).* Dot size reflects the percentage of cells expressing each gene, and color intensity represents the average expression level within each cluster

**Figure S6.**
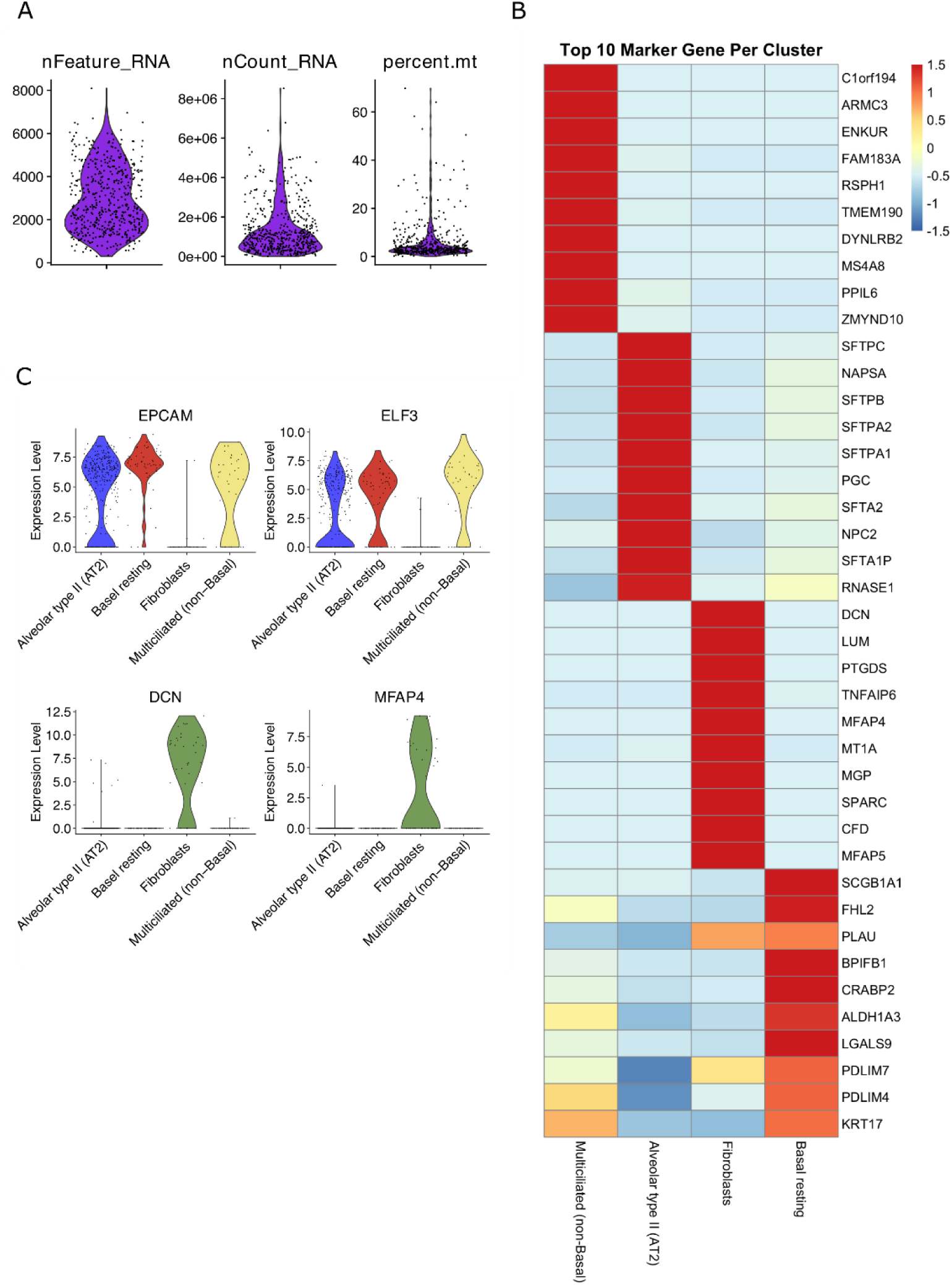
Single-cell RNA-seq quality control, clustering, and marker gene expression in lung samples. **(A)** Violin plots showing distributions of standard single-cell RNA-seq quality control metrics across all cells, including the number of detected genes (nFeature_RNA), total UMI counts (nCount_RNA), and the percentage of mitochondrial transcripts (percent.mt). **(B)** Heatmap of the top 10 differentially expressed genes per cluster, highlighting transcriptionally distinct epithelial and stromal compartments. Genes are ranked by average log-normalized expression within each cluster relative to others. Color scale indicates scaled expression levels. **(C)** Violin plots depicting normalized expression levels of representative markers: epithelial markers *EPCAM* and *ELF3* enriched in epithelial groups, and stromal/extracellular matrix markers *DCN* and *MFAP4* enriched in fibroblasts.

**Figure S7.**
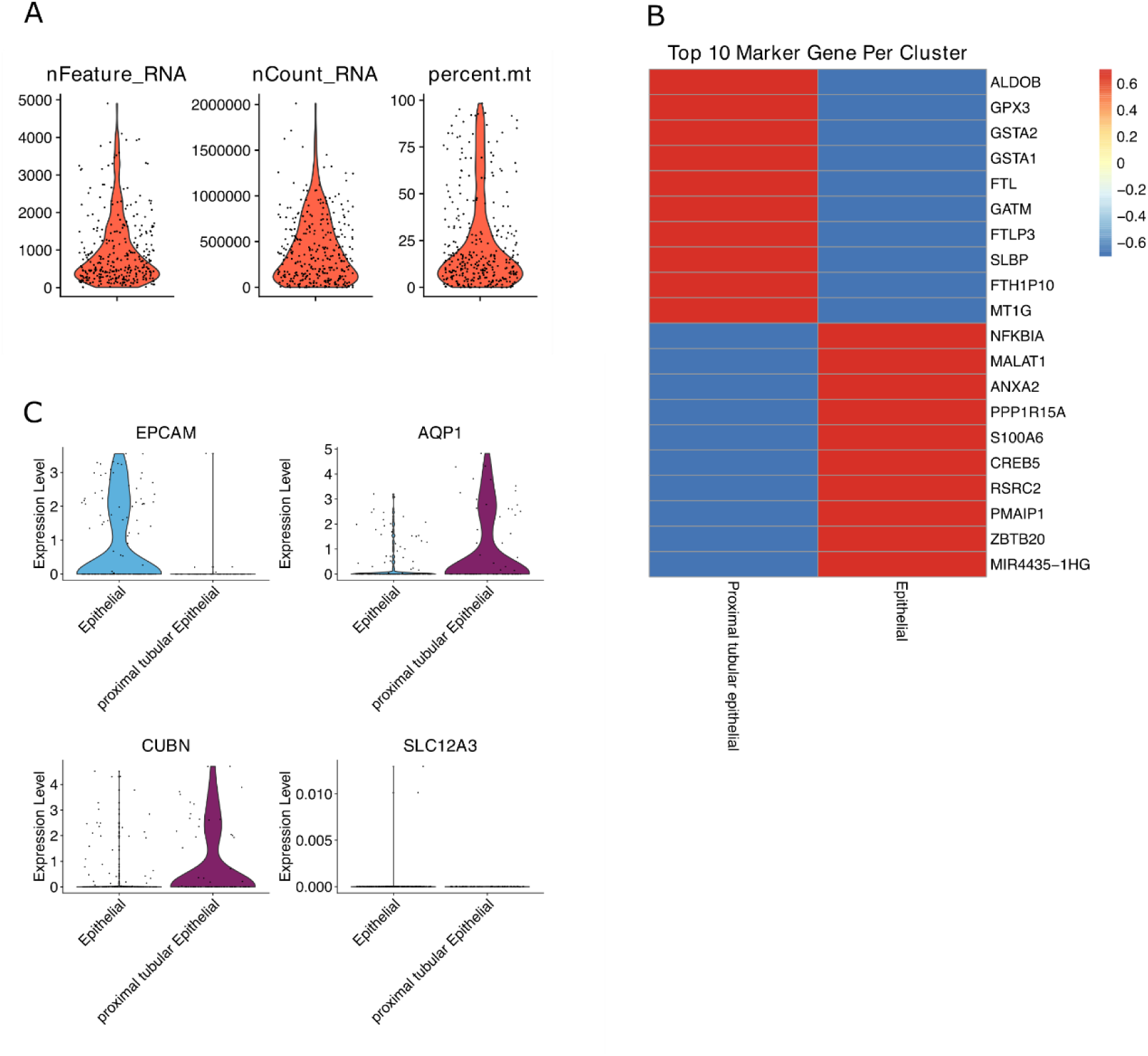
Single-cell RNA-seq quality control, clustering, and marker gene expression in renal samples. **(A)** Violin plots showing distributions of standard single-cell RNA-seq quality control metrics across all cells, including the number of detected genes (nFeature_RNA), total UMI counts (nCount_RNA), and the percentage of mitochondrial transcripts (percent.mt). **(B)** Heatmap of the top 10 differentially expressed genes per cluster, separating proximal tubular epithelial cells from broader epithelial populations. Genes are ranked by average log-normalized expression within each cluster relative to others. Color scale indicates scaled expression levels. **(C)** Violin plots depicting normalized expression levels of representative markers: epithelial marker *EPCAM* enriched in epithelial cells, proximal-tubule markers *AQP1* and *CUBN* enriched in the proximal tubular epithelial cluster and *SLC12A3*, expressed in the distal convoluted tubule.

**Figure S8.**
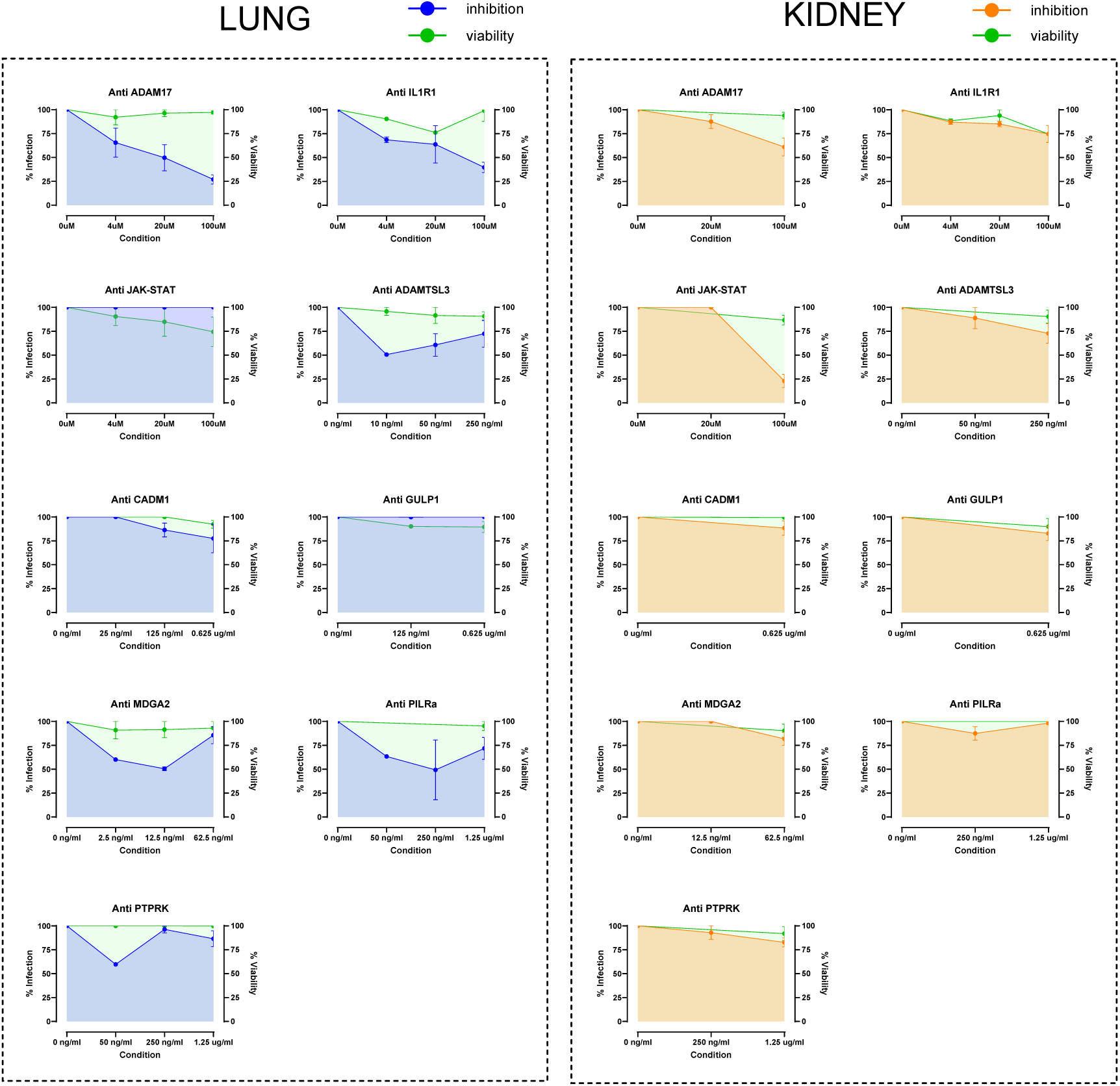
Dose-response and cell-viability profiles for drug treatments in lung and kidney cell models under SARS-CoV-2 pseudovirus infection. Dose-response curves showing target inhibition (blue for lung, orange for kidney) and corresponding cell viability (green for both) following treatment with increasing concentrations of the indicated inhibitor. Each panel represents an individual inhibitor tested in either lung (left) or kidney (right) cell models. Inhibition and viability values were normalized to untreated controls and are presented as mean ± s.e.m. from n=1-3 biological replicates. Shaded areas denote the 95% confidence interval of the fitted response curves.

**Figure S9.**
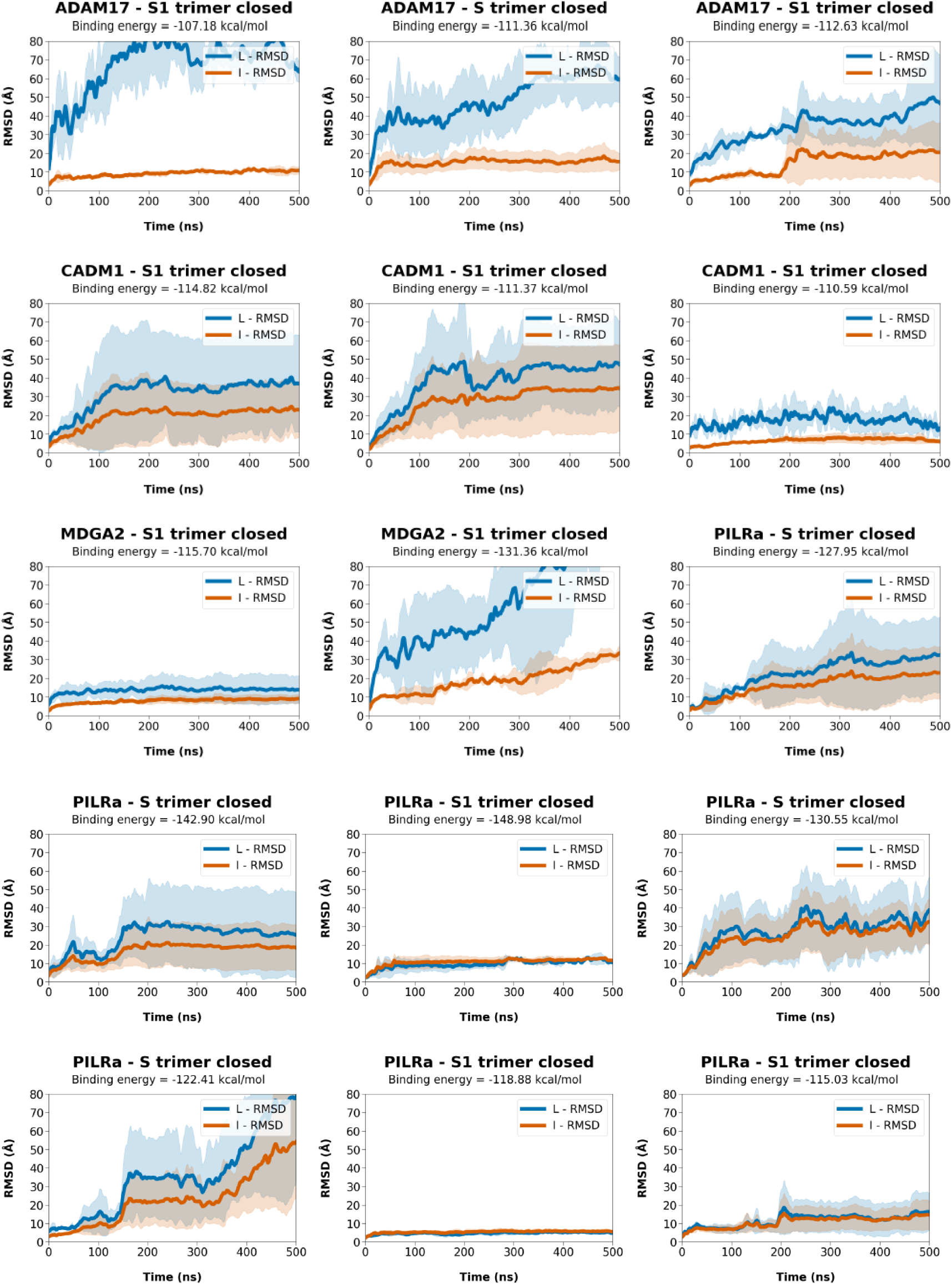
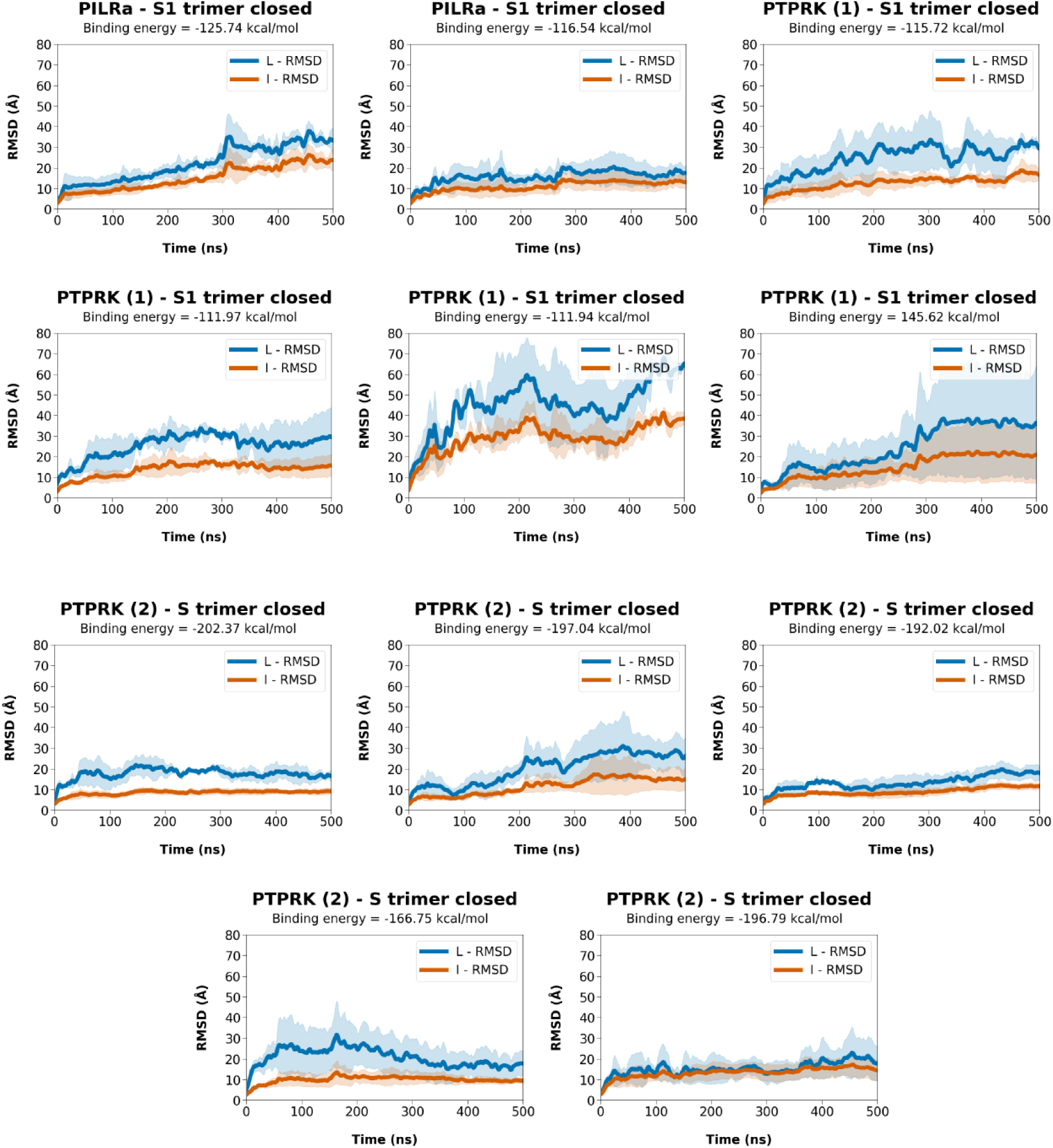
Molecular dynamics simulation results of predicted poses with highest interaction affinities. Each curve represents the average RMSD metric obtained from three independent replicas, after smoothing. The shaded area indicates the standard deviation among replicas.

**Table S2.**
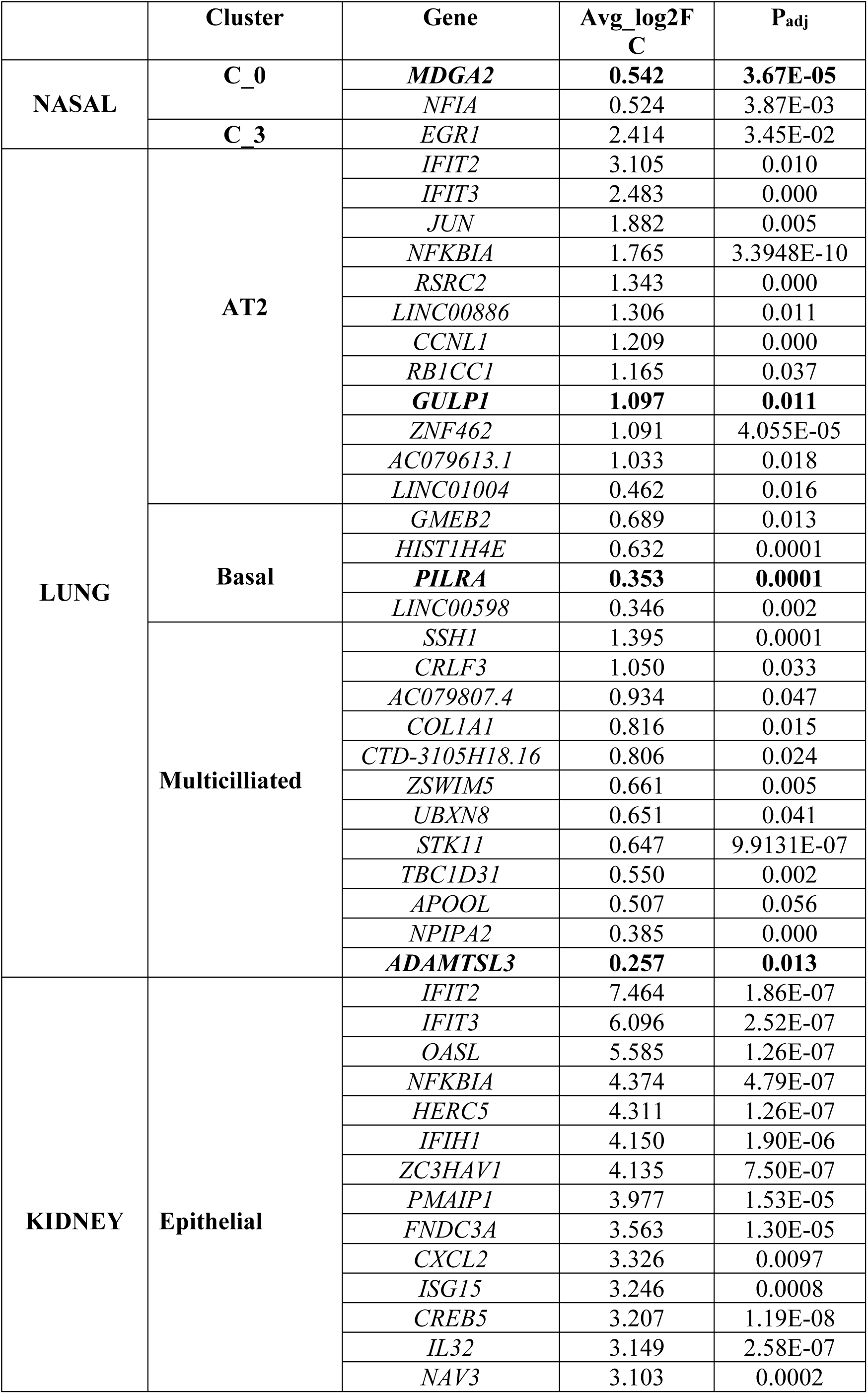
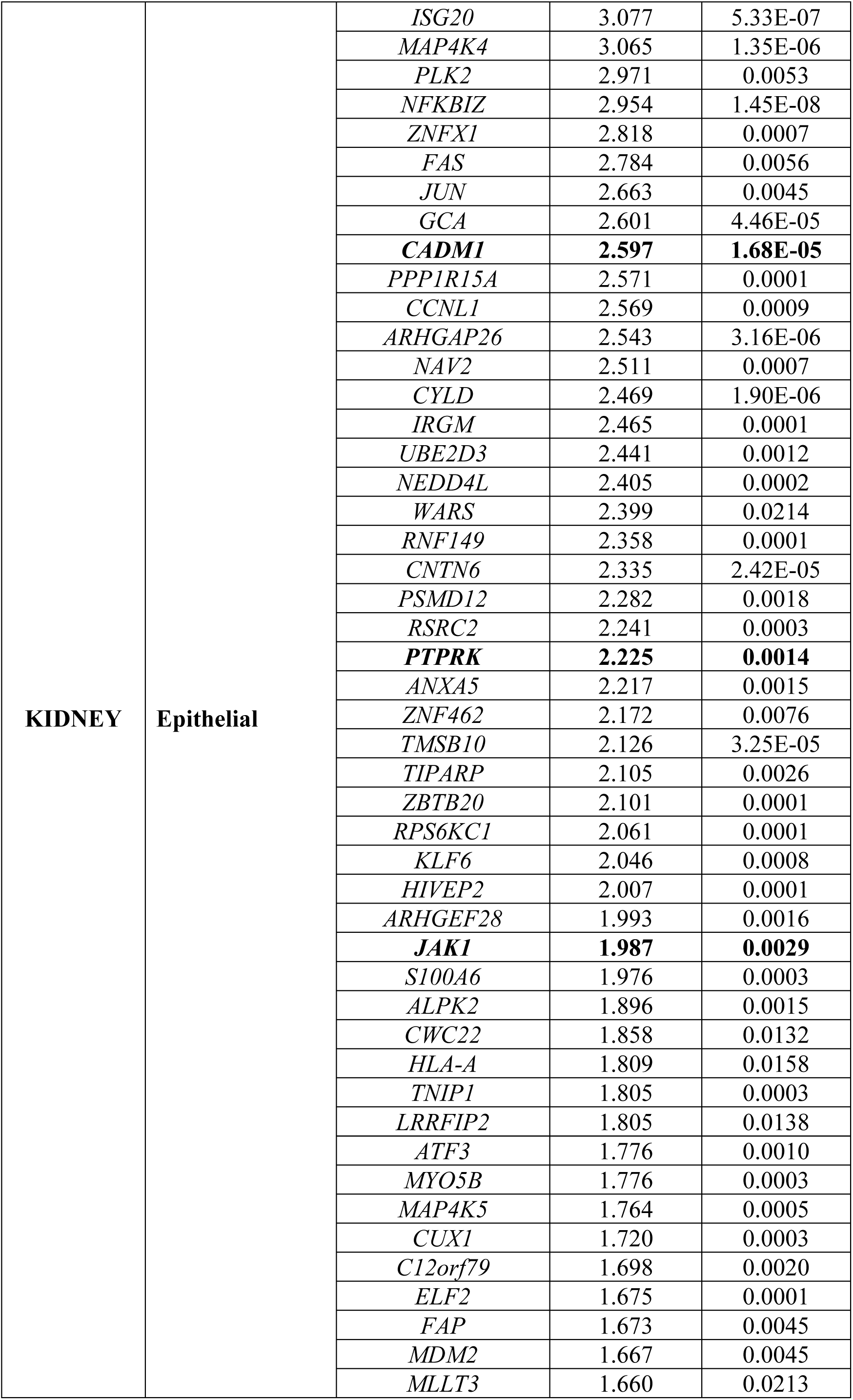
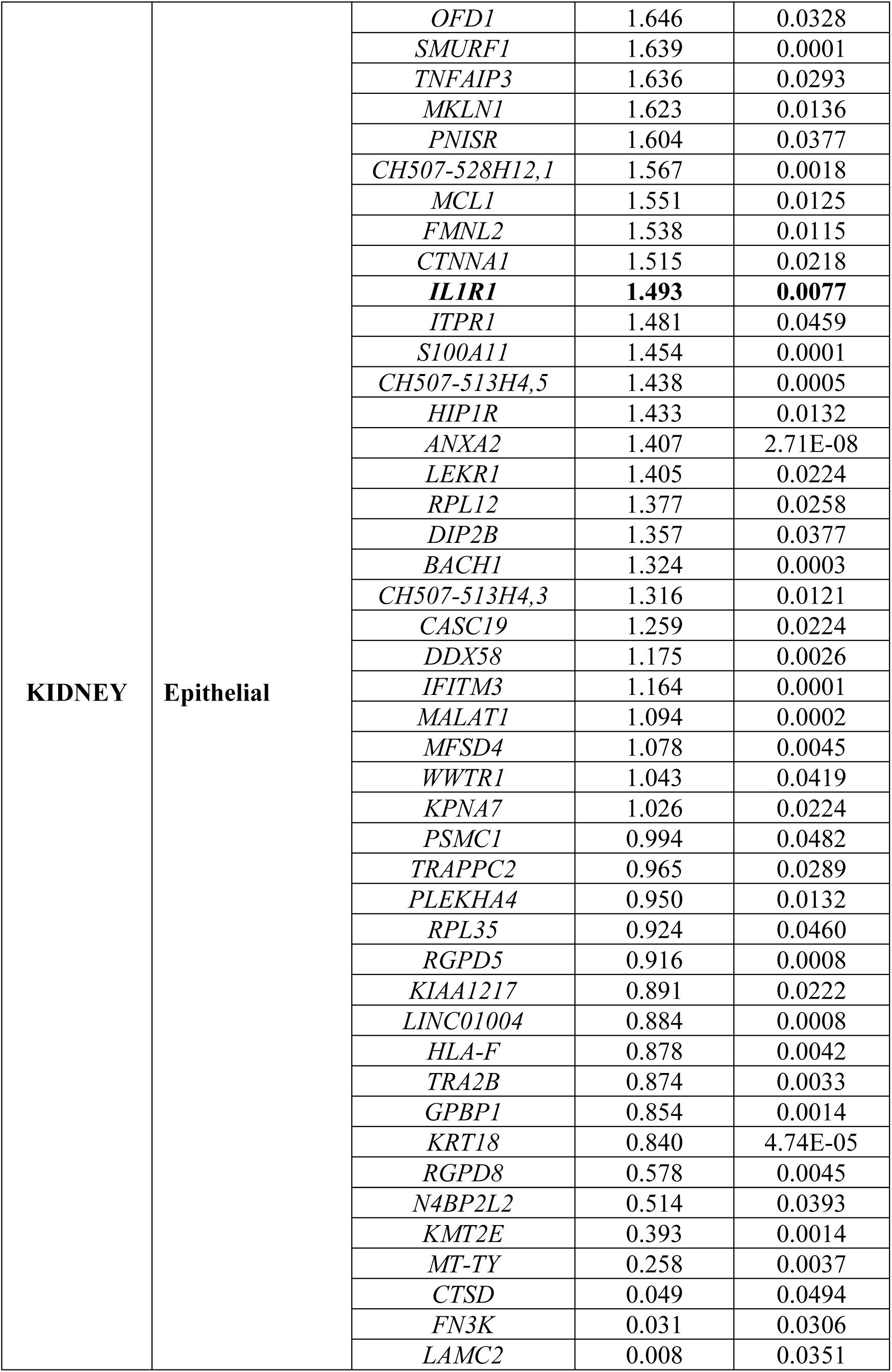
Upregulated genes by tissue-cluster.

**Table S3.** Median gene expression of reported entry factors and phenotypic markers by GFP expression and tissue cluster.

| GENE |  | NASAL |  |  |  |  | LUNG |  |  |  | KIDNEY |  |
| --- | --- | --- | --- | --- | --- | --- | --- | --- | --- | --- | --- | --- |
|  |  | C_0 | C_1 | C_2 | C_3 | C_4 | AT2 | Basal | Fibrob. | Multicil. | Epithel. | PTEC |
| ACE2 | GFP+ | 0.000 | 0.000 | 0.000 | 0.000 | 0.000 | 0.061 | 0.000 | 0.000 | 0.000 | 0.096 | 0.008 |
|  | GFP- | 0.000 | 0.000 | 0.000 | 0.000 | 0.000 | 0.034 | 0.036 | 0.000 | 0.000 | 0.013 | 0.259 |
| ADAM10 | GFP+ | 0.215 | 0.516 | 0.146 | 0.425 | 0.027 | 0.670 | 2.103 | 0.566 | 0.728 | 0.055 | 0.061 |
|  | GFP- | 0.000 | 0.736 | 0.150 | 0.012 | 0.423 | 0.823 | 2.172 | 0.795 | 1.088 | 0.001 | 0.101 |
| ADAM17 | GFP+ | 0.227 | 0.042 | 0.241 | 0.171 | 0.037 | 0.702 | 1.210 | 0.510 | 0.000 | 0.070 | 0.043 |
|  | GFP- | 0.004 | 0.192 | 0.240 | 0.382 | 0.221 | 0.776 | 1.348 | 1.166 | 1.340 | 0.057 | 0.000 |
| ADAMTSL3 | GFP+ | 0.031 | 0.120 | 0.000 | 0.083 | 0.000 | 0.143 | 0.355 | 0.189 | 0.627 | 0.076 | 0.100 |
|  | GFP- | 0.133 | 0.101 | 0.023 | 0.000 | 0.000 | 0.175 | 0.197 | 0.000 | 0.138 | 0.107 | 0.000 |
| ANPEP | GFP+ | 0.000 | 0.002 | 0.000 | 0.039 | 0.000 | 0.000 | 0.660 | 0.000 | 0.494 | 0.132 | 0.286 |
|  | GFP- | 0.088 | 0.008 | 0.000 | 0.000 | 0.000 | 0.042 | 0.079 | 0.000 | 0.000 | 0.344 | 0.295 |
| ASGR1 | GFP+ | 0.062 | 0.000 | 0.000 | 0.000 | 0.000 | 0.024 | 0.238 | 0.000 | 0.000 | 0.000 | 0.000 |
|  | GFP- | 0.106 | 0.000 | 0.000 | 0.002 | 0.004 | 0.144 | 0.261 | 0.000 | 0.000 | 0.000 | 0.000 |
| BSG | GFP+ | 0.137 | 0.301 | 0.117 | 0.215 | 0.000 | 2.737 | 4.107 | 2.658 | 2.503 | 0.823 | 0.352 |
|  | GFP- | 0.120 | 0.654 | 0.143 | 0.027 | 0.275 | 3.904 | 5.078 | 1.812 | 3.612 | 0.542 | 0.598 |
| CADM1 | GFP+ | 0.249 | 0.055 | 0.076 | 0.125 | 0.100 | 2.598 | 0.412 | 0.066 | 0.000 | 1.046 | 0.065 |
|  | GFP- | 0.568 | 0.402 | 0.217 | 0.345 | 0.372 | 2.829 | 1.323 | 0.756 | 1.207 | 0.000 | 0.001 |
| CD14 | GFP+ | 0.017 | 0.000 | 0.000 | 0.008 | 0.000 | 0.016 | 0.404 | 0.264 | 0.000 | 0.000 | 0.000 |
|  | GFP- | 0.000 | 0.000 | 0.000 | 0.000 | 0.000 | 0.041 | 0.000 | 0.000 | 0.000 | 0.000 | 0.000 |
| CD209 | GFP+ | 0.000 | 0.000 | 0.000 | 0.000 | 0.000 | 0.000 | 0.262 | 0.000 | 0.000 | 0.000 | 0.000 |
|  | GFP- | 0.000 | 0.000 | 0.000 | 0.000 | 0.000 | 0.000 | 0.000 | 0.000 | 0.000 | 0.000 | 0.000 |
| CDH1 | GFP+ | 0.252 | 0.465 | 0.521 | 0.293 | 0.008 | 1.755 | 2.667 | 0.000 | 2.941 | 0.097 | 0.117 |
|  | GFP- | 0.401 | 0.519 | 0.407 | 0.611 | 0.311 | 2.345 | 4.163 | 0.218 | 2.707 | 0.057 | 0.000 |
| CLIC1 | GFP+ | 0.191 | 1.106 | 0.248 | 0.473 | 0.000 | 4.353 | 7.409 | 2.150 | 6.750 | 1.144 | 0.301 |
|  | GFP- | 0.017 | 1.506 | 0.000 | 0.214 | 0.233 | 4.860 | 7.293 | 2.794 | 6.226 | 0.479 | 0.000 |
| CTSL | GFP+ | 0.026 | 0.177 | 0.000 | 0.000 | 0.000 | 0.919 | 1.522 | 3.589 | 2.252 | 0.399 | 0.201 |
|  | GFP- | 0.000 | 0.035 | 0.000 | 0.220 | 0.000 | 1.247 | 2.181 | 5.047 | 0.767 | 0.254 | 0.796 |
| DPP4 | GFP+ | 0.000 | 0.000 | 0.000 | 0.000 | 0.000 | 0.838 | 0.039 | 0.000 | 0.437 | 0.353 | 0.386 |
|  | GFP- | 0.000 | 0.000 | 0.000 | 0.000 | 0.000 | 0.902 | 0.249 | 0.000 | 0.064 | 0.219 | 0.272 |
| EPCAM | GFP+ | 0.137 | 0.844 | 0.207 | 0.283 | 0.000 | 4.189 | 6.035 | 0.268 | 4.889 | 0.981 | 0.094 |
|  | GFP- | 0.174 | 0.862 | 0.269 | 0.128 | 0.175 | 4.783 | 6.900 | 0.063 | 4.836 | 0.549 | 0.000 |
| FURIN | GFP+ | 0.000 | 0.004 | 0.073 | 0.000 | 0.000 | 0.197 | 0.270 | 0.000 | 0.000 | 0.001 | 0.000 |
|  | GFP- | 0.000 | 0.000 | 0.063 | 0.000 | 0.000 | 0.237 | 0.028 | 0.687 | 0.039 | 0.000 | 0.042 |
| GULP1 | GFP+ | 0.674 | 0.762 | 0.620 | 0.000 | 0.332 | 1.874 | 2.060 | 0.626 | 4.089 | 0.514 | 0.115 |
|  | GFP- | 0.330 | 0.626 | 0.752 | 0.084 | 0.102 | 0.788 | 2.267 | 1.260 | 3.383 | 0.021 | 0.000 |
| HLA-DRA | GFP+ | 0.161 | 0.242 | 0.000 | 0.341 | 0.000 | 4.472 | 2.424 | 0.978 | 2.636 | 0.241 | 0.245 |
|  | GFP- | 0.002 | 0.005 | 0.000 | 0.141 | 0.188 | 4.308 | 4.530 | 0.752 | 4.529 | 0.073 | 0.111 |
| IL1R1 | GFP+ | 0.000 | 0.144 | 0.006 | 0.093 | 0.000 | 1.351 | 1.373 | 4.123 | 0.000 | 0.500 | 0.047 |
|  | GFP- | 0.002 | 0.179 | 0.007 | 0.223 | 0.000 | 1.010 | 1.848 | 3.768 | 0.355 | 0.000 | 0.000 |
| ITGA6 | GFP+ | 0.211 | 0.200 | 0.035 | 0.012 | 0.749 | 0.442 | 0.884 | 0.000 | 0.000 | 0.104 | 0.059 |
|  | GFP- | 0.002 | 1.016 | 0.075 | 0.065 | 0.022 | 0.799 | 1.379 | 0.000 | 0.516 | 0.225 | 0.053 |
| JAK1 | GFP+ | 0.315 | 0.592 | 0.389 | 0.710 | 0.000 | 2.828 | 3.761 | 1.603 | 3.624 | 0.999 | 0.248 |
|  | GFP- | 0.456 | 0.396 | 0.392 | 0.455 | 0.652 | 2.714 | 3.676 | 2.057 | 2.199 | 0.099 | 0.008 |
| JAK2 | GFP+ | 0.075 | 0.241 | 0.002 | 0.000 | 0.000 | 0.541 | 0.908 | 1.083 | 2.261 | 0.133 | 0.001 |
|  | GFP- | 0.003 | 0.128 | 0.032 | 0.000 | 0.000 | 0.309 | 0.794 | 0.000 | 0.924 | 0.057 | 0.000 |
| KREMEN1 | GFP+ | 0.002 | 0.000 | 0.025 | 0.000 | 0.000 | 0.063 | 0.452 | 0.245 | 0.560 | 0.003 | 0.075 |
|  | GFP- | 0.053 | 0.000 | 0.018 | 0.004 | 0.006 | 0.061 | 0.103 | 0.000 | 0.184 | 0.000 | 0.018 |
| MDGA2 | GFP+ | 0.085 | 0.000 | 0.002 | 0.000 | 0.000 | 0.127 | 0.154 | 0.051 | 0.139 | 0.032 | 0.106 |
|  | GFP- | 0.006 | 0.000 | 0.010 | 0.003 | 0.000 | 0.101 | 0.100 | 0.326 | 0.234 | 0.037 | 0.000 |
| MME | GFP+ | 0.002 | 0.002 | 0.084 | 0.001 | 0.000 | 0.104 | 0.275 | 0.460 | 0.110 | 0.117 | 0.602 |
|  | GFP- | 0.046 | 0.002 | 0.000 | 0.000 | 0.001 | 0.134 | 0.059 | 0.603 | 0.099 | 0.457 | 0.517 |
| MUC5AC | GFP+ | 0.000 | 0.000 | 0.000 | 0.001 | 0.000 | 0.003 | 0.321 | 0.000 | 0.000 | 0.000 | 0.000 |
|  | GFP- | 0.070 | 0.000 | 0.000 | 0.000 | 0.000 | 0.057 | 0.018 | 0.000 | 0.000 | 0.000 | 0.000 |
| NGFR | GFP+ | 0.069 | 0.000 | 0.002 | 0.000 | 0.208 | 0.003 | 0.297 | 0.000 | 0.000 | 0.000 | 0.000 |
|  | GFP- | 0.091 | 0.000 | 0.000 | 0.216 | 0.015 | 0.000 | 0.000 | 0.000 | 0.000 | 0.000 | 0.000 |
| NPM1 | GFP+ | 1.051 | 2.452 | 0.469 | 2.636 | 1.325 | 5.230 | 7.292 | 4.457 | 6.618 | 1.246 | 0.286 |
|  | GFP- | 0.205 | 2.586 | 0.685 | 1.185 | 1.814 | 6.230 | 7.717 | 4.557 | 4.918 | 0.772 | 0.283 |
| NRP1 | GFP+ | 0.134 | 0.074 | 0.157 | 0.000 | 0.000 | 0.480 | 1.701 | 0.324 | 0.561 | 0.097 | 0.225 |
|  | GFP- | 0.006 | 0.053 | 0.000 | 0.000 | 0.000 | 1.057 | 1.562 | 0.699 | 0.857 | 0.006 | 0.000 |
| PILRα | GFP+ | 0.026 | 0.033 | 0.027 | 0.000 | 0.000 | 0.049 | 0.362 | 0.380 | 0.000 | 0.039 | 0.000 |
|  | GFP- | 0.289 | 0.000 | 0.000 | 0.000 | 0.002 | 0.036 | 0.178 | 0.000 | 0.000 | 0.000 | 0.000 |
| PODXL | GFP+ | 0.076 | 0.027 | 0.000 | 0.039 | 0.000 | 0.499 | 0.854 | 0.444 | 0.979 | 0.121 | 0.000 |
|  | GFP- | 0.000 | 0.091 | 0.000 | 0.211 | 0.034 | 0.214 | 0.501 | 0.000 | 0.244 | 0.000 | 0.004 |
| PTPRK | GFP+ | 1.302 | 0.851 | 0.510 | 1.001 | 3.133 | 2.815 | 4.546 | 0.532 | 3.537 | 0.784 | 0.050 |
|  | GFP- | 1.677 | 0.922 | 1.048 | 0.474 | 1.467 | 2.764 | 4.713 | 0.505 | 3.673 | 0.001 | 0.017 |
| SFTPC | GFP+ | 0.000 | 0.000 | 0.000 | 0.000 | 0.000 | 9.889 | 2.594 | 0.673 | 0.161 | 0.000 | 0.000 |
|  | GFP- | 0.000 | 0.000 | 0.000 | 0.000 | 0.000 | 9.283 | 2.475 | 0.000 | 0.494 | 0.000 | 0.000 |
| SLC1A5 | GFP+ | 0.149 | 0.136 | 0.029 | 0.095 | 0.000 | 0.977 | 0.303 | 0.660 | 0.728 | 0.116 | 0.000 |
|  | GFP- | 0.124 | 0.102 | 0.020 | 0.000 | 0.000 | 1.541 | 2.181 | 1.585 | 0.670 | 0.037 | 0.000 |
| TMEM106B | GFP+ | 0.230 | 0.375 | 0.000 | 0.000 | 0.000 | 0.797 | 1.669 | 0.189 | 0.543 | 0.035 | 0.125 |
|  | GFP- | 0.000 | 0.289 | 0.286 | 0.000 | 0.000 | 0.893 | 2.041 | 0.000 | 1.976 | 0.098 | 0.000 |
| TMPRSS2 | GFP+ | 0.120 | 0.113 | 0.231 | 0.005 | 0.000 | 0.538 | 0.573 | 0.000 | 0.000 | 0.055 | 0.000 |
|  | GFP- | 0.019 | 0.108 | 0.000 | 0.001 | 0.019 | 0.451 | 0.637 | 0.000 | 0.690 | 0.022 | 0.000 |
| TUBA1A | GFP+ | 0.014 | 0.116 | 0.002 | 0.028 | 0.000 | 1.305 | 3.967 | 1.385 | 4.291 | 0.155 | 0.088 |
|  | GFP- | 0.013 | 0.220 | 0.000 | 0.249 | 0.114 | 2.339 | 4.151 | 2.999 | 2.466 | 0.049 | 0.043 |

**Table S4.**
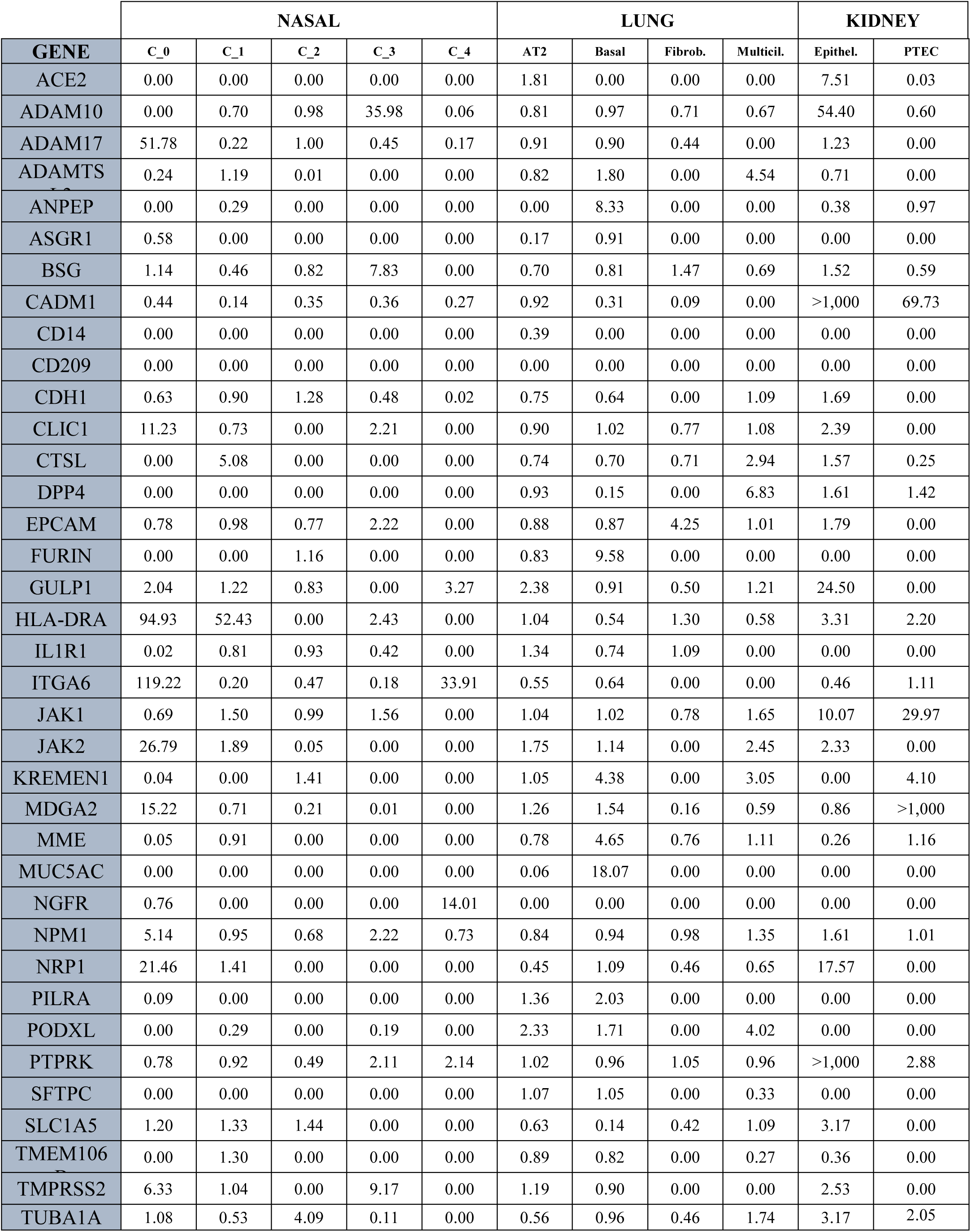
Ratio of median expression (from table S1) in GFP^+^ vs GFP^-^.

## REFERENCES

1. Delorey, T.M., Ziegler, C.G.K., Heimberg, G., Normand, R., Yang, Y., Segerstolpe, Å., Abbondanza, D., Fleming, S.J., Subramanian, A., Montoro, D.T., et al. (2021). COVID-19 tissue atlases reveal SARS-CoV-2 pathology and cellular targets. Nature 595, 107–113. 10.1038/s41586-021-03570-8.

2. Puelles, V.G., Lütgehetmann, M., Lindenmeyer, M.T., Sperhake, J.P., Wong, M.N., Allweiss, L., Chilla, S., Heinemann, A., Wanner, N., Liu, S., et al. (2020). Multiorgan and Renal Tropism of SARS-CoV-2. N. Engl. J. Med. 383, 590–592. 10.1056/NEJMc2011400.

3. Jackson, C.B., Farzan, M., Chen, B., and Choe, H. (2022). Mechanisms of SARS-CoV-2 entry into cells. Nat. Rev. Mol. Cell Biol. 23, 3–20. 10.1038/s41580-021-00418-x.

4. Jansen, J., Reimer, K.C., Nagai, J.S., Varghese, F.S., Overheul, G.J., de Beer, M., Roverts, R., Daviran, D., Fermin, L.A.S., Willemsen, B., et al. (2022). SARS-CoV-2 infects the human kidney and drives fibrosis in kidney organoids. Cell Stem Cell 29, 217–231.e8. 10.1016/j.stem.2021.12.010.

5. Hoffmann, M., Kleine-Weber, H., Schroeder, S., Krüger, N., Herrler, T., Erichsen, S., Schiergens, T.S., Herrler, G., Wu, N.-H., Nitsche, A., et al. (2020). SARS-CoV-2 Cell Entry Depends on ACE2 and TMPRSS2 and Is Blocked by a Clinically Proven Protease Inhibitor. Cell 181, 271–280.e8. 10.1016/j.cell.2020.02.052.

6. Shang, J., Wan, Y., Luo, C., Ye, G., Geng, Q., Auerbach, A., and Li, F. (2020). Cell entry mechanisms of SARS-CoV-2. Proc. Natl. Acad. Sci. 117, 11727–11734. 10.1073/pnas.2003138117.

7. Grau-Expósito, J., Perea, D., Suppi, M., Massana, N., Vergara, A., Soler, M.J., Trinite, B., Blanco, J., García-Pérez, J., Alcamí, J., et al. (2022). Evaluation of SARS-CoV-2 entry, inflammation and new therapeutics in human lung tissue cells. PLOS Pathog. 18, e1010171. 10.1371/journal.ppat.1010171.

8. HCA Lung Biological Network, Sungnak, W., Huang, N., Bécavin, C., Berg, M., Queen, R., Litvinukova, M., Talavera-López, C., Maatz, H., Reichart, D., et al. (2020). SARS-CoV-2 entry factors are highly expressed in nasal epithelial cells together with innate immune genes. Nat. Med. 26, 681–687. 10.1038/s41591-020-0868-6.

9. Ziegler, C.G.K., Allon, S.J., Nyquist, S.K., Mbano, I.M., Miao, V.N., Tzouanas, C.N., Cao, Y., Yousif, A.S., Bals, J., Hauser, B.M., et al. (2020). SARS-CoV-2 Receptor ACE2 Is an Interferon-Stimulated Gene in Human Airway Epithelial Cells and Is Detected in Specific Cell Subsets across Tissues. Cell 181, 1016–1035.e19. 10.1016/j.cell.2020.04.035.

10. Lukassen, S., Chua, R.L., Trefzer, T., Kahn, N.C., Schneider, M.A., Muley, T., Winter, H., Meister, M., Veith, C., Boots, A.W., et al. (2020). SARS -CoV-2 receptor ACE 2 and TMPRSS 2 are primarily expressed in bronchial transient secretory cells. EMBO J. 39. 10.15252/embj.20105114.

11. Muus, C., Luecken, M.D., Eraslan, G., Sikkema, L., Waghray, A., Heimberg, G., Kobayashi, Y., Vaishnav, E.D., Subramanian, A., Smillie, C., et al. (2021). Single-cell meta-analysis of SARS-CoV-2 entry genes across tissues and demographics. Nat. Med. 27, 546–559. 10.1038/s41591-020-01227-z.

12. Ahn, J.H., Kim, J., Hong, S.P., Choi, S.Y., Yang, M.J., Ju, Y.S., Kim, Y.T., Kim, H.M., Rahman, M.T., Chung, M.K., et al. (2021). Nasal ciliated cells are primary targets for SARS-CoV-2 replication in the early stage of COVID-19. J. Clin. Invest. 131. 10.1172/JCI148517.

13. Zhang, H., Li, H.-B., Lyu, J.-R., Lei, X.-M., Li, W., Wu, G., Lyu, J., and Dai, Z.-M. (2020). Specific ACE2 expression in small intestinal enterocytes may cause gastrointestinal symptoms and injury after 2019-nCoV infection. Int. J. Infect. Dis. 96, 19–24. 10.1016/j.ijid.2020.04.027.

14. Carossino, M., Izadmehr, S., Trujillo, J.D., Gaudreault, N.N., Dittmar, W., Morozov, I., Balasuriya, U.B.R., Cordon-Cardo, C., García-Sastre, A., and Richt, J.A. (2024). ACE2 and TMPRSS2 distribution in the respiratory tract of different animal species and its correlation with SARS-CoV-2 tissue tropism. Microbiol. Spectr. 12. 10.1128/spectrum.03270-23.

15. Evans, J.P., and Liu, S.-L. (2021). Role of host factors in SARS-CoV-2 entry. J. Biol. Chem. 297, 100847. 10.1016/j.jbc.2021.100847.

16. Baggen, J., Vanstreels, E., Jansen, S., and Daelemans, D. (2021). Cellular host factors for SARS-CoV-2 infection. Nat. Microbiol. 6, 1219–1232. 10.1038/s41564-021-00958-0.

17. Cantuti-Castelvetri, L., Ojha, R., Pedro, L.D., Djannatian, M., Franz, J., Kuivanen, S., van der Meer, F., Kallio, K., Kaya, T., Anastasina, M., et al. (2020). Neuropilin-1 facilitates SARS-CoV-2 cell entry and infectivity. Science 370, 856–860. 10.1126/science.abd2985.

18. Lambert, D.W., Yarski, M., Warner, F.J., Thornhill, P., Parkin, E.T., Smith, A.I., Hooper, N.M., and Turner, A.J. (2005). Tumor Necrosis Factor-α Convertase (ADAM17) Mediates Regulated Ectodomain Shedding of the Severe-acute Respiratory Syndrome-Coronavirus (SARS-CoV) Receptor, Angiotensin-converting Enzyme-2 (ACE2). J. Biol. Chem. 280, 30113–30119. 10.1074/jbc.M505111200.

19. Behboudi, E., Nooreddin Faraji, S., Daryabor, G., Mohammad Ali Hashemi, S., Asadi, M., Edalat, F., Javad Raee, M., and Hatam, G. (2024). SARS-CoV-2 mechanisms of cell tropism in various organs considering host factors. Heliyon 10, e26577. 10.1016/j.heliyon.2024.e26577.

20. Jocher, G., Grass, V., Tschirner, S.K., Riepler, L., Breimann, S., Kaya, T., Oelsner, M., Hamad, M.S., Hofmann, L.I., Blobel, C.P., et al. (2022). ADAM10 and ADAM17 promote SARS-CoV-2 cell entry and spike protein-mediated lung cell fusion. EMBO Rep. 23. 10.15252/embr.202154305.

21. Yeung, M.L., Teng, J.L.L., Jia, L., Zhang, C., Huang, C., Cai, J.-P., Zhou, R., Chan, K.-H., Zhao, H., Zhu, L., et al. (2021). Soluble ACE2-mediated cell entry of SARS-CoV-2 via interaction with proteins related to the renin-angiotensin system. Cell 184, 2212–2228.e12. 10.1016/j.cell.2021.02.053.

22. Walls, A.C., Park, Y.-J., Tortorici, M.A., Wall, A., McGuire, A.T., and Veesler, D. (2020). Structure, Function, and Antigenicity of the SARS-CoV-2 Spike Glycoprotein. Cell 181, 281–292.e6. 10.1016/j.cell.2020.02.058.

23. Wang, K., Chen, W., Zhang, Z., Deng, Y., Lian, J.-Q., Du, P., Wei, D., Zhang, Y., Sun, X.-X., Gong, L., et al. (2020). CD147-spike protein is a novel route for SARS-CoV-2 infection to host cells. Signal Transduct. Target. Ther. 5. 10.1038/s41392-020-00426-x.

24. Gu, Y., Cao, J., Zhang, X., Gao, H., Wang, Y., Wang, J., He, J., Jiang, X., Zhang, J., Shen, G., et al. (2022). Receptome profiling identifies KREMEN1 and ASGR1 as alternative functional receptors of SARS-CoV-2. Cell Res. 32, 24–37. 10.1038/s41422-021-00595-6.

25. Baggen, J., Jacquemyn, M., Persoons, L., Vanstreels, E., Pye, V.E., Wrobel, A.G., Calvaresi, V., Martin, S.R., Roustan, C., Cronin, N.B., et al. (2023). TMEM106B is a receptor mediating ACE2-independent SARS-CoV-2 cell entry. Cell 186, 3427–3442.e22. 10.1016/j.cell.2023.06.005.

26. Singh, M., Bansal, V., and Feschotte, C. (2020). A Single-Cell RNA Expression Map of Human Coronavirus Entry Factors. Cell Rep. 32. 10.1016/j.celrep.2020.108175.

27. Sunshine, S., Puschnik, A.S., Replogle, J.M., Laurie, M.T., Liu, J., Zha, B.S., Nuñez, J.K., Byrum, J.R., McMorrow, A.H., Frieman, M.B., et al. (2023). Systematic functional interrogation of SARS-CoV-2 host factors using Perturb-seq. Nat. Commun. 14. 10.1038/s41467-023-41788-4.

28. Le Pen, J., Paniccia, G., Kinast, V., Moncada-Velez, M., Ashbrook, A.W., Bauer, M., Hoffmann, H.-H., Pinharanda, A., Ricardo-Lax, I., Stenzel, A.F., et al. (2024). A genome-wide arrayed CRISPR screen identifies PLSCR1 as an intrinsic barrier to SARS-CoV-2 entry that recent virus variants have evolved to resist. PLOS Biol. 22, e3002767. 10.1371/journal.pbio.3002767.

29. Hou, J., Wei, Y., Zou, J., Jaffery, R., Sun, L., Liang, S., Zheng, N., Guerrero, A.M., Egan, N.A., Bohat, R., et al. (2024). Integrated multi-omics analyses identify anti-viral host factors and pathways controlling SARS-CoV-2 infection. Nat. Commun. 15. 10.1038/s41467-023-44175-1.

30. Daly, J.L., Simonetti, B., Klein, K., Chen, K.-E., Williamson, M.K., Antón-Plágaro, C., Shoemark, D.K., Simón-Gracia, L., Bauer, M., Hollandi, R., et al. (2020). Neuropilin-1 is a host factor for SARS-CoV-2 infection. Science 370, 861–865. 10.1126/science.abd3072.

31. Verma, S., Patel, C.N., and Chandra, M. (2021). Identification of novel inhibitors of SARS-COV -2 main protease (M ^pro^ ) from *Withania* sp. by molecular docking and molecular dynamics simulation. J. Comput. Chem. 42, 1861–1872. 10.1002/jcc.26717.

32. Nagavarapu, S., Kumar, J., and Biswal, P.K. (2025). Molecular docking based comparative study of antiviral compounds on SARS-CoV-2 spike protein. Nat. Prod. Res. 39, 4975–4983. 10.1080/14786419.2024.2355589.

33. Bestion, E., Halfon, P., Mezouar, S., and Mège, J.-L. (2022). Cell and Animal Models for SARS-CoV-2 Research. Viruses 14, 1507. 10.3390/v14071507.

34. Khundmiri, S.J., Chen, L., Lederer, E.D., Yang, C.-R., and Knepper, M.A. (2021). Transcriptomes of Major Proximal Tubule Cell Culture Models. J. Am. Soc. Nephrol. 32, 86–97. 10.1681/ASN.2020010009.

35. Steiner, S., Kratzel, A., Barut, G.T., Lang, R.M., Aguiar Moreira, E., Thomann, L., Kelly, J.N., and Thiel, V. (2024). SARS-CoV-2 biology and host interactions. Nat. Rev. Microbiol. 22, 206–225. 10.1038/s41579-023-01003-z.

36. Peng, L., Gao, L., Wu, X., Fan, Y., Liu, M., Chen, J., Song, J., Kong, J., Dong, Y., Li, B., et al. (2022). Lung Organoids as Model to Study SARS-CoV-2 Infection. Cells 11, 2758. 10.3390/cells11172758.

37. Mykytyn, A.Z., Breugem, T.I., Geurts, M.H., Beumer, J., Schipper, D., van Acker, R., van den Doel, P.B., van Royen, M.E., Zhang, J., Clevers, H., et al. (2023). SARS-CoV-2 Omicron entry is type II transmembrane serine protease-mediated in human airway and intestinal organoid models. J. Virol. 97. 10.1128/jvi.00851-23.

38. Mykytyn, A.Z., Breugem, T.I., Riesebosch, S., Schipper, D., van den Doel, P.B., Rottier, R.J., Lamers, M.M., and Haagmans, B.L. (2021). SARS-CoV-2 entry into human airway organoids is serine protease-mediated and facilitated by the multibasic cleavage site. eLife 10. 10.7554/eLife.64508.

39. Vanslambrouck, J.M., Neil, J.A., Rudraraju, R., Mah, S., Tan, K.S., Groenewegen, E., Forbes, T.A., Karavendzas, K., Elliott, D.A., Porrello, E.R., et al. (2024). Kidney organoids reveal redundancy in viral entry pathways during ACE2-dependent SARS-CoV-2 infection. J. Virol. 98. 10.1128/jvi.01802-23.

40. Woodall, M.N.J., Cujba, A.-M., Worlock, K.B., Case, K.-M., Masonou, T., Yoshida, M., Polanski, K., Huang, N., Lindeboom, R.G.H., Mamanova, L., et al. (2024). Age-specific nasal epithelial responses to SARS-CoV-2 infection. Nat. Microbiol. 9, 1293–1311. 10.1038/s41564-024-01658-1.

41. Hicks, W., Hall, L., Sigurdson, L., Stewart, C., Hard, R., Winston, J., and Lwebuga-Mukasa, J. (1997). Isolation and Characterization of Basal Cells from Human Upper Respiratory Epithelium. Exp. Cell Res. 237, 357–363. 10.1006/excr.1997.3796.

42. Hewitt, R.J., and Lloyd, C.M. (2021). Regulation of immune responses by the airway epithelial cell landscape. Nat. Rev. Immunol. 21, 347–362. 10.1038/s41577-020-00477-9.

43. Bonser, L.R., Koh, K.D., Johansson, K., Choksi, S.P., Cheng, D., Liu, L., Sun, D.I., Zlock, L.T., Eckalbar, W.L., Finkbeiner, W.E., et al. (2021). Flow-Cytometric Analysis and Purification of Airway Epithelial-Cell Subsets. Am. J. Respir. Cell Mol. Biol. 64, 308–317. 10.1165/rcmb.2020-0149MA.

44. Goerlich, N., Brand, H.A., Langhans, V., Tesch, S., Schachtner, T., Koch, B., Paliege, A., Schneider, W., Grützkau, A., Reinke, P., et al. (2020). Kidney transplant monitoring by urinary flow cytometry: Biomarker combination of T cells, renal tubular epithelial cells, and podocalyxin-positive cells detects rejection. Sci. Rep. 10. 10.1038/s41598-020-57524-7.

45. Jackson, C.L., and Bottier, M. (2022). Methods for the assessment of human airway ciliary function. Eur. Respir. J. 60. 10.1183/13993003.02300-2021.

46. Bocciolini, C., Nappi, E., Giunta, G., Paoletti, G., Malvezzi, L., Monti, G., Macchi, A., Amorosa, L., and Heffler, E. (2023). Middle meatus nasal cytology compared to inferior turbinate cytology in non allergic rhinitis. Eur. Arch. Otorhinolaryngol. 280, 913–918. 10.1007/s00405-022-07629-8.

47. Chapman, H.A., Li, X., Alexander, J.P., Brumwell, A., Lorizio, W., Tan, K., Sonnenberg, A., Wei, Y., and Vu, T.H. (2011). Integrin α6β4 identifies an adult distal lung epithelial population with regenerative potential in mice. J. Clin. Invest. 121, 2855–2862. 10.1172/JCI57673.

48. Wang, Y., Huang, Z., Nayak, P.S., Matthews, B.D., Warburton, D., Shi, W., and Sanchez-Esteban, J. (2013). Strain-induced differentiation of fetal type II epithelial cells is mediated via the integrin α6β1-ADAM17/tumor necrosis factor-α-converting enzyme (TACE) signaling pathway. J. Biol. Chem. 288, 25646–25657. 10.1074/jbc.M113.473777.

49. Trzpis, M., Popa, E.R., McLaughlin, P.M.J., van Goor, H., Timmer, A., Bosman, G.W., de Leij, L.M.F.H., and Harmsen, M.C. (2007). Spatial and Temporal Expression Patterns of the Epithelial Cell Adhesion Molecule (EpCAM/EGP-2) in Developing and Adult Kidneys. Nephron Exp. Nephrol. 107, e119–e131. 10.1159/000111039.

50. Picelli, S., Faridani, O.R., Björklund, Å.K., Winberg, G., Sagasser, S., and Sandberg, R. (2014). Full-length RNA-seq from single cells using Smart-seq2. Nat. Protoc. 9, 171–181. 10.1038/nprot.2014.006.

51. Sikkema, L., Ramírez-Suástegui, C., Strobl, D.C., Gillett, T.E., Zappia, L., Madissoon, E., Markov, N.S., Zaragosi, L.-E., Ji, Y., Ansari, M., et al. (2023). An integrated cell atlas of the lung in health and disease. Nat. Med. 29, 1563–1577. 10.1038/s41591-023-02327-2.

52. Namestnikov, M., Cohen-Zontag, O., Omer, D., Gnatek, Y., Goldberg, S., Vincent, T., Singh, S., Shiber, Y., Rafaeli Yehudai, T., Volkov, H., et al. (2025). Human fetal kidney organoids model early human nephrogenesis and Notch-driven cell fate. EMBO J. 44, 4681–4719. 10.1038/s44318-025-00504-2.

53. Kim, S., Jang, G., Kim, H., Lim, D., Han, K.A., Um, J.W., and Ko, J. (2024). MDGAs perform activity-dependent synapse type-specific suppression via distinct extracellular mechanisms. Proc. Natl. Acad. Sci. 121. 10.1073/pnas.2322978121.

54. Chau, D.D.-L., Yu, Z., Chan, W.W.R., Yuqi, Z., Chang, R.C.C., Ngo, J.C.K., Chan, H.Y.E., and Lau, K.-F. (2024). The cellular adaptor GULP1 interacts with ATG14 to potentiate autophagy and APP processing. Cell. Mol. Life Sci. 81. 10.1007/s00018-024-05351-8.

55. Rypdal, K.B., Olav Melleby, A., Robinson, E.L., Li, J., Palmero, S., Seifert, D.E., Martin, D., Clark, C., López, B., Andreassen, K., et al. (2022). ADAMTSL3 knock-out mice develop cardiac dysfunction and dilatation with increased TGFβ signalling after pressure overload. Commun. Biol. 5. 10.1038/s42003-022-04361-1.

56. Satoh, T., Arii, J., Suenaga, T., Wang, J., Kogure, A., Uehori, J., Arase, N., Shiratori, I., Tanaka, S., Kawaguchi, Y., et al. (2008). PILRα Is a Herpes Simplex Virus-1 Entry Coreceptor That Associates with Glycoprotein B. Cell 132, 935–944. 10.1016/j.cell.2008.01.043.

57. Haga, S., Yamamoto, N., Nakai-Murakami, C., Osawa, Y., Tokunaga, K., Sata, T., Yamamoto, N., Sasazuki, T., and Ishizaka, Y. (2008). Modulation of TNF-α-converting enzyme by the spike protein of SARS-CoV and ACE2 induces TNF-α production and facilitates viral entry. Proc. Natl. Acad. Sci. 105, 7809–7814. 10.1073/pnas.0711241105.

58. Scheller, J., Chalaris, A., Garbers, C., and Rose-John, S. (2011). ADAM17: a molecular switch to control inflammation and tissue regeneration. Trends Immunol. 32, 380–387. 10.1016/j.it.2011.05.005.

59. Franzetti, M., Forastieri, A., Borsa, N., Pandolfo, A., Molteni, C., Borghesi, L., Pontiggia, S., Evasi, G., Guiotto, L., Erba, M., et al. (2021). IL-1 Receptor Antagonist Anakinra in the Treatment of COVID-19 Acute Respiratory Distress Syndrome: A Retrospective, Observational Study. J. Immunol. 206, 1569–1575. 10.4049/jimmunol.2001126.

60. Nakazawa, D., Takeda, Y., Kanda, M., Tomaru, U., Ogawa, H., Kudo, T., Shiratori-Aso, S., Watanabe-Kusunoki, K., Ueda, Y., Miyoshi, A., et al. (2023). Inhibition of Toll-like receptor 4 and Interleukin-1 receptor prevent SARS-CoV-2 mediated kidney injury. Cell Death Discov. 9. 10.1038/s41420-023-01584-x.

61. Chen, D.-Y., Khan, N., Close, B.J., Goel, R.K., Blum, B., Tavares, A.H., Kenney, D., Conway, H.L., Ewoldt, J.K., Chitalia, V.C., et al. (2021). SARS-CoV-2 Disrupts Proximal Elements in the JAK-STAT Pathway. J. Virol. 95. 10.1128/JVI.00862-21.

62. Jankowski, J., Lee, H.K., Wilflingseder, J., and Hennighausen, L. (2021). JAK inhibitors dampen activation of interferon-activated transcriptomes and the SARS-CoV-2 receptor ACE2 in human renal proximal tubules. iScience 24, 102928. 10.1016/j.isci.2021.102928.

63. Takeuchi, F., Sugano, A., Yoneshige, A., Hagiyama, M., Inoue, T., Wada, A., Takaoka, Y., and Ito, A. (2024). Potential Contribution of Cell Adhesion Molecule 1 to the Binding of SARS-CoV-2 Spike Protein to Mouse Nasal Mucosa. Cells Tissues Organs 213, 326–337. 10.1159/000534892.

64. Nain, Z., Rana, H.K., Liò, P., Islam, S.M.S., Summers, M.A., and Moni, M.A. (2021). Pathogenetic profiling of COVID-19 and SARS-like viruses. Brief. Bioinform. 22, 1175–1196. 10.1093/bib/bbaa173.

65. Nguyen, L., McCord, K.A., Bui, D.T., Bouwman, K.M., Kitova, E.N., Elaish, M., Kumawat, D., Daskhan, G.C., Tomris, I., Han, L., et al. (2022). Sialic acid-containing glycolipids mediate binding and viral entry of SARS-CoV-2. Nat. Chem. Biol. 18, 81–90. 10.1038/s41589-021-00924-1.

66. Korber, B., Fischer, W.M., Gnanakaran, S., Yoon, H., Theiler, J., Abfalterer, W., Hengartner, N., Giorgi, E.E., Bhattacharya, T., Foley, B., et al. (2020). Tracking Changes in SARS-CoV-2 Spike: Evidence that D614G Increases Infectivity of the COVID-19 Virus. Cell 182, 812–827.e19. 10.1016/j.cell.2020.06.043.

67. Shuai, H., Chan, J.F.-W., Hu, B., Chai, Y., Yuen, T.T.-T., Yin, F., Huang, X., Yoon, C., Hu, J.-C., Liu, H., et al. (2022). Attenuated replication and pathogenicity of SARS-CoV-2 B.1.1.529 Omicron. Nature 603, 693–699. 10.1038/s41586-022-04442-5.

68. Mungmunpuntipantip, R., and Wiwanitkit, V. (2022). Change in binding affinity with ACE2 receptor in beta, delta and omicron SARS CoV2 variants. Int. J. Physiol. Pathophysiol. Pharmacol. 14, 124–128.

69. Whitt, M.A. (2010). Generation of VSV pseudotypes using recombinant ΔG-VSV for studies on virus entry, identification of entry inhibitors, and immune responses to vaccines. J. Virol. Methods 169, 365–374. 10.1016/j.jviromet.2010.08.006.

70. Parekh, S., Ziegenhain, C., Vieth, B., Enard, W., and Hellmann, I. (2018). zUMIs - A fast and flexible pipeline to process RNA sequencing data with UMIs. GigaScience 7. 10.1093/gigascience/giy059.

71. Hao, Y., Stuart, T., Kowalski, M.H., Choudhary, S., Hoffman, P., Hartman, A., Srivastava, A., Molla, G., Madad, S., Fernandez-Granda, C., et al. (2024). Dictionary learning for integrative, multimodal and scalable single-cell analysis. Nat. Biotechnol. 42, 293–304. 10.1038/s41587-023-01767-y.

72. Yang, Z., Zeng, X., Zhao, Y., and Chen, R. (2023). AlphaFold2 and its applications in the fields of biology and medicine. Signal Transduct. Target. Ther. 8. 10.1038/s41392-023-01381-z.

73. Mirdita, M., Schütze, K., Moriwaki, Y., Heo, L., Ovchinnikov, S., and Steinegger, M. (2022). ColabFold: making protein folding accessible to all. Nat. Methods 19, 679–682. 10.1038/s41592-022-01488-1.

74. Waterhouse, A., Bertoni, M., Bienert, S., Studer, G., Tauriello, G., Gumienny, R., Heer, F.T., de Beer, T.A.P., Rempfer, C., Bordoli, L., et al. (2018). SWISS-MODEL: homology modelling of protein structures and complexes. Nucleic Acids Res. 46, W296–W303. 10.1093/nar/gky427.

75. Cheng, T.M.-K., Blundell, T.L., and Fernandez-Recio, J. (2007). pyDock: electrostatics and desolvation for effective scoring of rigid-body protein-protein docking. Proteins 68, 503–515. 10.1002/prot.21419.

76. Gainza, P., Sverrisson, F., Monti, F., Rodolà, E., Boscaini, D., Bronstein, M.M., and Correia, B.E. (2020). Deciphering interaction fingerprints from protein molecular surfaces using geometric deep learning. Nat. Methods 17, 184–192. 10.1038/s41592-019-0666-6.

77. Madhavi Sastry, G., Adzhigirey, M., Day, T., Annabhimoju, R., and Sherman, W. (2013). Protein and ligand preparation: parameters, protocols, and influence on virtual screening enrichments. J. Comput. Aided Mol. Des. 27, 221–234. 10.1007/s10822-013-9644-8.

78. Abramson, J., Adler, J., Dunger, J., Evans, R., Green, T., Pritzel, A., Ronneberger, O., Willmore, L., Ballard, A.J., Bambrick, J., et al. (2024). Accurate structure prediction of biomolecular interactions with AlphaFold 3. Nature 630, 493–500. 10.1038/s41586-024-07487-w.

79. Borrelli, K.W., Vitalis, A., Alcantara, R., and Guallar, V. (2005). PELE: Protein Energy Landscape Exploration. A Novel Monte Carlo Based Technique. J. Chem. Theory Comput. 1, 1304–1311. 10.1021/ct0501811.

80. Metropolis, N., Rosenbluth, A.W., Rosenbluth, M.N., Teller, A.H., and Teller, E. (1953). Equation of State Calculations by Fast Computing Machines. J. Chem. Phys. 21, 1087–1092. 10.1063/1.1699114.

81. Pronk, S., Páll, S., Schulz, R., Larsson, P., Bjelkmar, P., Apostolov, R., Shirts, M.R., Smith, J.C., Kasson, P.M., van der Spoel, D., et al. (2013). GROMACS 4.5: a high-throughput and highly parallel open source molecular simulation toolkit. Bioinforma. Oxf. Engl. 29, 845–854. 10.1093/bioinformatics/btt055.

82. Huang, J., and MacKerell, A.D. (2013). CHARMM36 all-atom additive protein force field: Validation based on comparison to NMR data. J. Comput. Chem. 34, 2135–2145. 10.1002/jcc.23354.

83. Mark, P., and Nilsson, L. (2001). Structure and Dynamics of the TIP3P, SPC, and SPC/E Water Models at 298 K. J. Phys. Chem. A 105, 9954–9960. 10.1021/jp003020w.

